# Evolutionary analysis implicates RNA polymerase II pausing and chromatin structure in nematode piRNA biogenesis

**DOI:** 10.1101/281360

**Authors:** T. Beltran, C. Barroso, T.Y. Birkle, L. Stevens, H. T. Schwartz, P.W. Sternberg, H. Fradin, K. Gunsalus, F. Piano, J. Cotton, E. Martínez-Pérez, M. Blaxter, P. Sarkies

## Abstract

Piwi-interacting RNAs (piRNAs) control transposable elements widely across metazoans but have rapidly evolving biogenesis pathways. In *Caenorhabditis elegans,* almost all piRNA loci are found within two 3Mb clusters on Chromosome IV. Each piRNA locus possesses an upstream motif that recruits RNA polymerase II to produce a ∼28 nt precursor transcript. Here, we use comparative epigenomics across nematodes to gain insight into piRNA biogenesis. We show that the piRNA upstream motif is derived from core promoter elements controlling snRNA biogenesis. We describe two alternative modes of piRNA organisation in nematodes: in *C. elegans* and closely related nematodes, piRNAs are clustered within repressive H3K27me3 chromatin, whilst in other species, typified by *Pristionchus pacificus,* piRNAs are distributed genome-wide within introns of actively transcribed genes. In both groups, piRNA production depends on downstream sequence signals associated with RNA polymerase II pausing, which synergise with the chromatin environment to control piRNA precursor transcription.

## Introduction

piRNAs are 21-30 nucleotide small RNAs that bind to members of the Piwi subfamily of Argonaute proteins. Conserved across animals, their ancestral role appears to be to defend the genome against transposable elements (TEs). piRNAs hybridise to TE-derived RNAs, instigating post-transcriptional and transcriptional silencing of TEs (Rödelsperger et al., 2017; Siomi et al., 2011). In many organisms, piRNAs are essential for fertility, and germ cell development is defective in their absence (Weick and Miska, 2014). piRNA biogenesis has been characterised in arthropods and mammals. In these taxa long precursor RNAs are produced from ∼100 genomic loci and processed into mature 26-30 nt piRNAs bound to Piwi proteins. In germ cells, primary piRNA processing occurs as part of a ping-pong cycle that amplifies piRNAs targeted to expressed TEs (Brennecke et al., 2007).

Despite the conservation of the general logic of piRNA pathway function and biogenesis across animals, the piRNA machinery diverges rapidly amongst closely related organisms. Many of the dedicated piRNA biogenesis factors characterised in *Drosophila melanogaster* are not conserved even amongst Diptera, let alone across arthropods (Weick and Miska, 2014). Some aspects of piRNA function are idiosyncratic. For example in the mosquito *Aedes aegypti* piRNAs derived from RNA viruses are found in the gut (Miesen et al., 2015) while in the silkworm *Bombyx mori* piRNAs regulate a specific protein-coding target gene in the sex determination pathway (Kiuchi et al., 2014). How and why piRNA function and biogenesis diverges so rapidly remains unclear.

Nematodes (phylum Nematoda) represent an extreme example of the diversity of the piRNA system. Although piRNAs are conserved within the Rhabditina (Clade V (Blaxter et al., 1998)), the entire piRNA pathway has been lost independently in several lineages across the phylum (Sarkies et al., 2015). Rhabditina contains the model nematode *Caenorhabditis elegans,* in which characterisation of piRNAs is most advanced. As in other metazoans, piRNAs in *C. elegans* (termed 21U, for their typical length and 5′ uracil) associate with a Piwi protein, PRG-1, target transposable elements and are important for fertility (Batista et al., 2008; Das et al., 2008). However, the *C. elegans* piRNA system displays intriguing differences to other organisms. First, the ping-pong cycle is not present. Instead, piRNAs recruit RNA dependent RNA polymerases (RdRPs) to target RNAs. RdRP activity produces 22 nt RNAs that have a 5′ G (22G RNAs), antisense to targets, which bind to nematode-specific argonautes (WAGO) to bring about silencing (Bagijn et al., 2012). Second, piRNAs are produced from monocistronic genomic loci, the vast majority of which are preceded by a GTTTC consensus motif (the Ruby motif) (Batista et al., 2008; Das et al., 2008; Ruby et al., 2006) that recruits RNA polymerase II (Pol II) to produce a 5′ capped 28 nucleotide precursor transcript (Billi et al., 2013; Gu et al., 2012). Two piRNA clusters on chromosome IV, spanning 2.5 Mb and 3.7 Mb, contain over 90% of piRNA loci (Ruby et al., 2006). Pol II transcription of piRNA precursors requires the nematode-specific pseudokinase PRDE-1 (Billi et al., 2013; Gu et al., 2012; Weick et al., 2014) and the small nuclear RNA (snRNA) activating protein complex (SNAPc) component SNPC-4 (Kasper et al., 2014). Together, SNPC-4 and PRDE-1 have been hypothesised to establish a specific chromatin structure essential for piRNA biogenesis (Kasper et al., 2014).

Several key aspects of nematode piRNA production remain poorly understood. Amongst the outstanding problems are the evolutionary origin of the Ruby motif, how Pol II is controlled such that it transcribes only very short piRNA precursors and how the clustering of piRNA loci into genomic regions contributes to piRNA production. Here, through a comparative analysis of piRNA biogenesis across nematodes, we obtain new insights into these questions. First, we uncover the evolutionary origin of the Ruby motif, showing that nematode piRNA transcription evolved from snRNA transcription. Second, we reveal two distinct modes of piRNA locus organisation in nematodes - either forming high density clusters within H3K27me3 chromatin domains or dispersed throughout the genome within active genes enriched for H3K36me3 chromatin. Third, we discover a novel downstream sequence signal at nematode piRNA loci, which likely promotes RNA Pol II pausing at piRNA loci to generate short piRNA precursors. By using CRISPR-mediated genome editing we confirm that both the surrounding chromatin environment and RNA Pol pausing sequence signals determine the activity of piRNA loci in nematodes.

## Results

### The Ruby motif is an ancient piRNA regulatory module

We surveyed a wide taxonomic span of nematodes for components of the piRNA system. All of the nematodes we examined within Rhabditina (Clade V) possessed the Piwi protein PRG-1 (Figure 1), with the exception of *Caenorhabditis plicata,* suggesting a recent loss of the piRNA pathway in this species. Outside of Clade V we only identified Piwi in the closely related free living nematodes *Plectus sambesii* and *Plectus murrayi.* The presence of Piwi in *P. sambesii,* which is ancestral to Rhabditida (which includes Clades III, IV and V), is consistent with the hypothesis that the piRNA pathway was lost independently in Clade III and Clade IV (Figure 1) (Sarkies et al., 2015).

**Figure 1.**
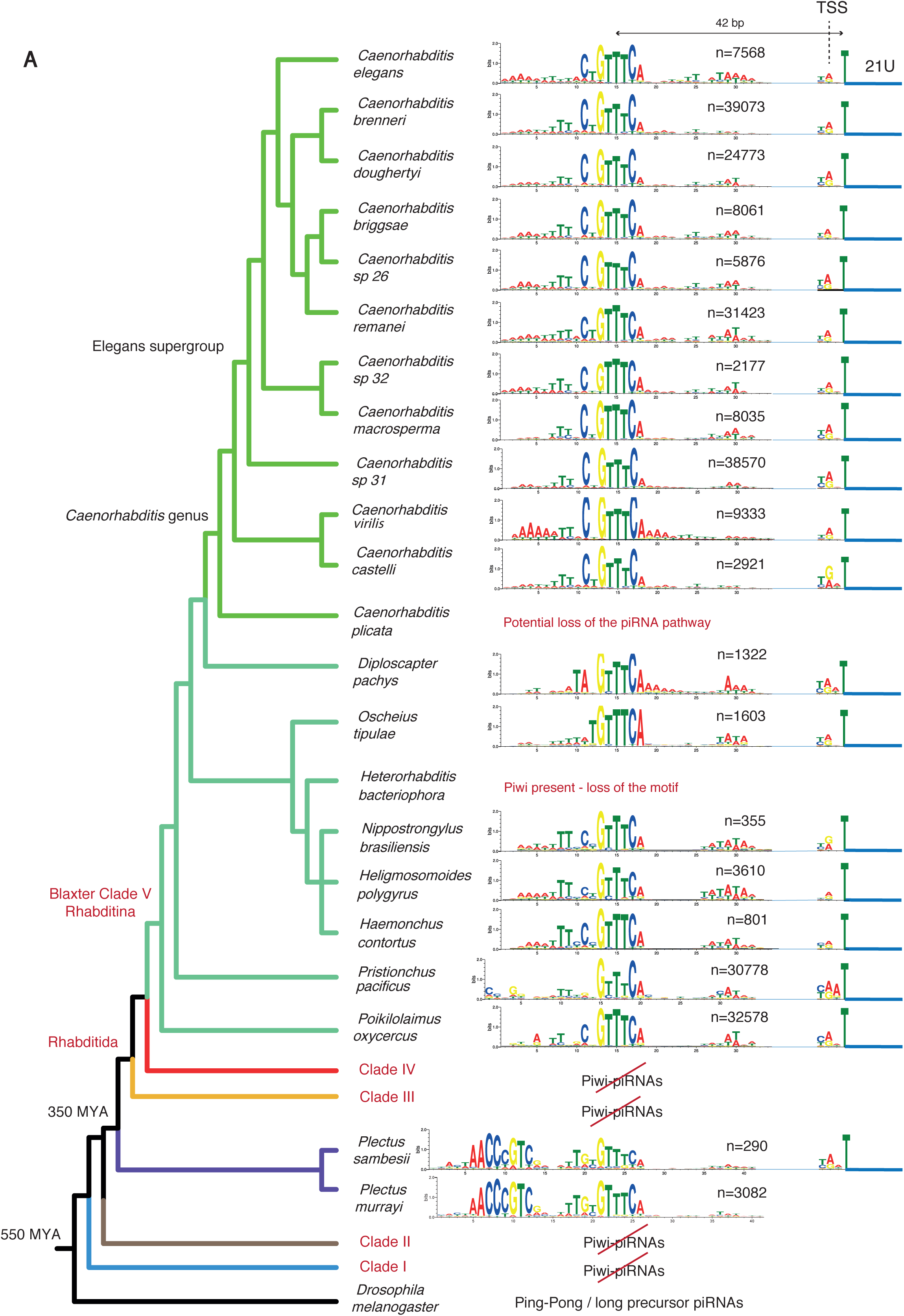
Overview of piRNA evolution in nematodes. Assesment of Piwi protein presence/absence, Ruby motif presence/absence, and number of piRNA loci with upstream motifs and unless otherwise indicated, 21U-RNA reads detected by high-throughput sequencing of small RNAs. Nematode phylogeny was taken from Koutsovolos and Blaxter, 2016.

To reveal the evolutionary dynamics of piRNA biogenesis we sequenced small RNAs and mapped them to their genomes in a number of additional species (Figure 1). We sequenced and assembled *de novo* the genomes of *P. sambesii* and *Poikilolaimus oxycercus* and sourced other genomes from ongoing and published projects. In the majority of nematodes with Piwi, including in *P. sambesii* and *P. murrayi,* we found a motif upstream of 21U-RNAs which bore remarkable similarity to the *C. elegans* core Ruby motif (Figure 1). We did not find any motifs upstream of any other type of small RNA. This suggests that the Ruby motif originated at least 350 mya (Figure 1). Surprisingly, there was no evidence of upstream motifs in *Heterorhabditis bacteriophora* despite the presence of Piwi, suggesting that that the Ruby motif has been lost in this species. In all nematodes with Ruby-motif dependent piRNAs transcription initiated two nucleotides upstream of the mature piRNA (Figure S1A), just as in *C. elegans* (Billi et al., 2013; Gu et al., 2012; Weick et al., 2014). We observed large variability in the number of piRNA loci across genomes (Figure 1) that could not be attributed to differences in sequencing depth. The biological and evolutionary correlates of these differences await future study.

### The Ruby motif is derived from the nematode SNAPc binding motif

The Ruby motif upstream of piRNAs in plectid nematodes was strongly associated with an additional 5′ motif (Figure 1A). The two motifs almost always occurred together (Figure S1B). This extra motif bears a striking similarity to the *C. elegans* SNAPc motifs, predicted from binding sites of the SNAPc subunit SNPC-4 (Kasper et al., 2014; Figure 2A). We predicted SNAPc motifs in *P. sambesii* from regions upstream of Pol II and Pol III-dependent snRNA genes. The *P. sambesii* SNAPc motif associated with snRNA genes and the piRNA motif are highly similar (Figure 2A). We recovered a TATA-box at Pol III-dependent snRNA loci, but not at Pol II snRNAs or piRNA loci, consistent with Pol II transcription of piRNAs (Hung and Stumph, 2011). We predicted SNAPc motifs in a wider sampling of nematodes with sequenced genomes, many of which have lost the piRNA pathway (Sarkies et al., 2015). Alignment of these motifs showed strong conservation of the 5′ half of the motif, whilst the 3′ half was more divergent. There were strong sequence similarities between the 3′ SNAPc motif and the Ruby motif in all nematodes (Figure S2-2), and the positioning of the 3′ SNAPc motif relative to snRNA transcription start sites (TSSs) mirrors the positioning of the Ruby motif at piRNA loci (Figure 2A). Comparison of SNAPc motifs from Clade II nematodes strongly suggested that the Ruby motif was derived from the 3′ half of an original nematode SNAPc motif and the two motifs diverged from each other since the last common ancestor of Rhabditida (Clades III, IV, V) (Figure 2, Figure S2-1). Altogether, our data suggests that nematode piRNA transcription evolved from snRNA transcription.

**Figure 2.**
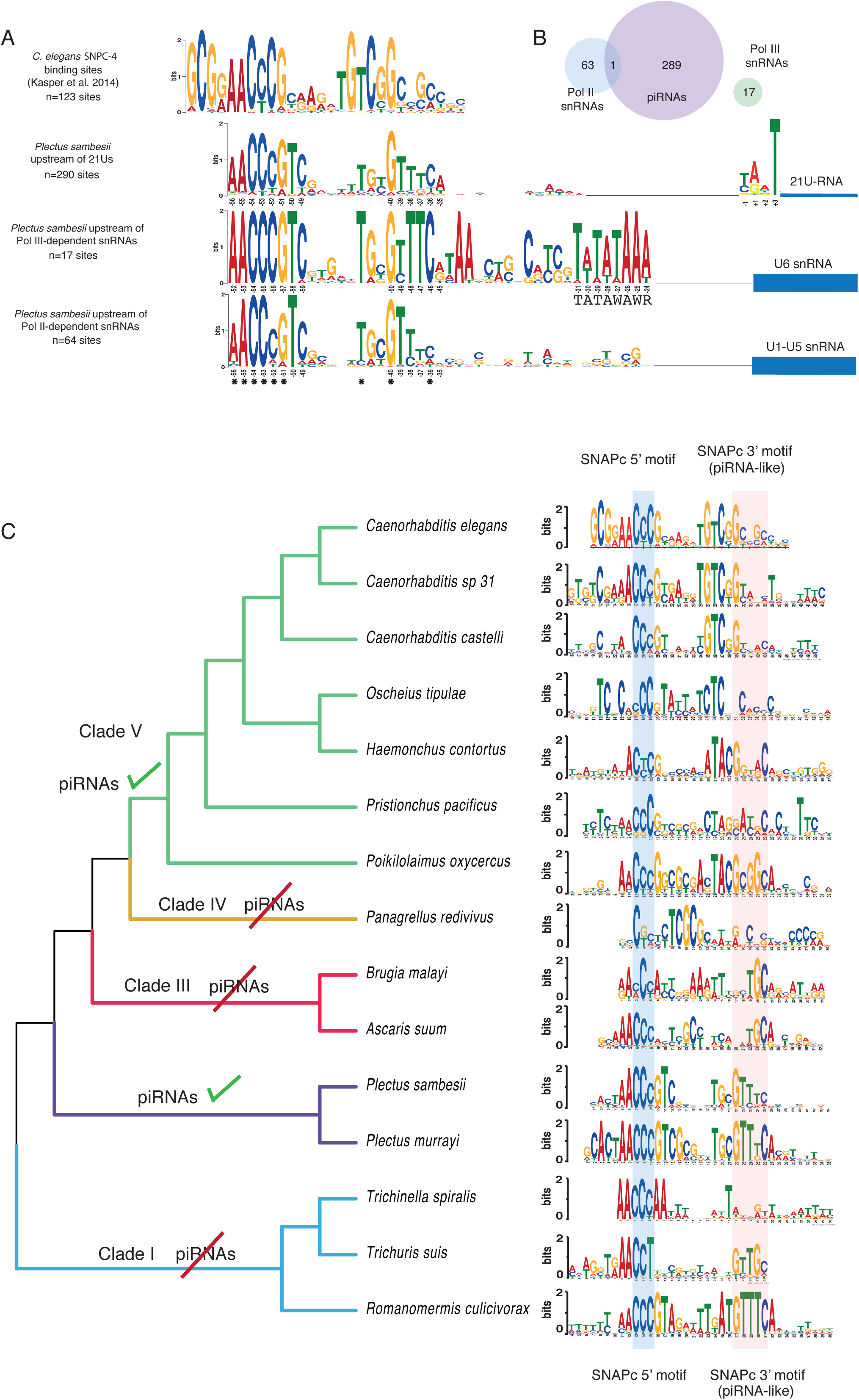
The nematode piRNA upstream motif evolved from the SNAPc binding motifs. A. Ruby and SNPC-4 motifs in *C. elegans* and *Plectus sambesii.* B. Evolution of SNPC-4 motifs defined upstream of annotated snRNA loci across nematodes with and without piRNAs.

### Two levels of piRNA clustering in nematodes

piRNA loci controlled by Ruby motifs in *C. elegans* are almost exclusively located in two ∼3 Mb regions on Chromosome IV. However, in the chromosomal genome assembly of *Pristionchus pacificus* (Rödelsperger et al., 2017), piRNA loci are distributed relatively evenly across the five autosomes but absent from the X chromosome (Figure 3A). Pristionchus is also in Rhabditina (Clade V) but is a member of Diplogasteromorpha, while *C. elegans* is in the Rhabditomorpha.

**Figure 3.**
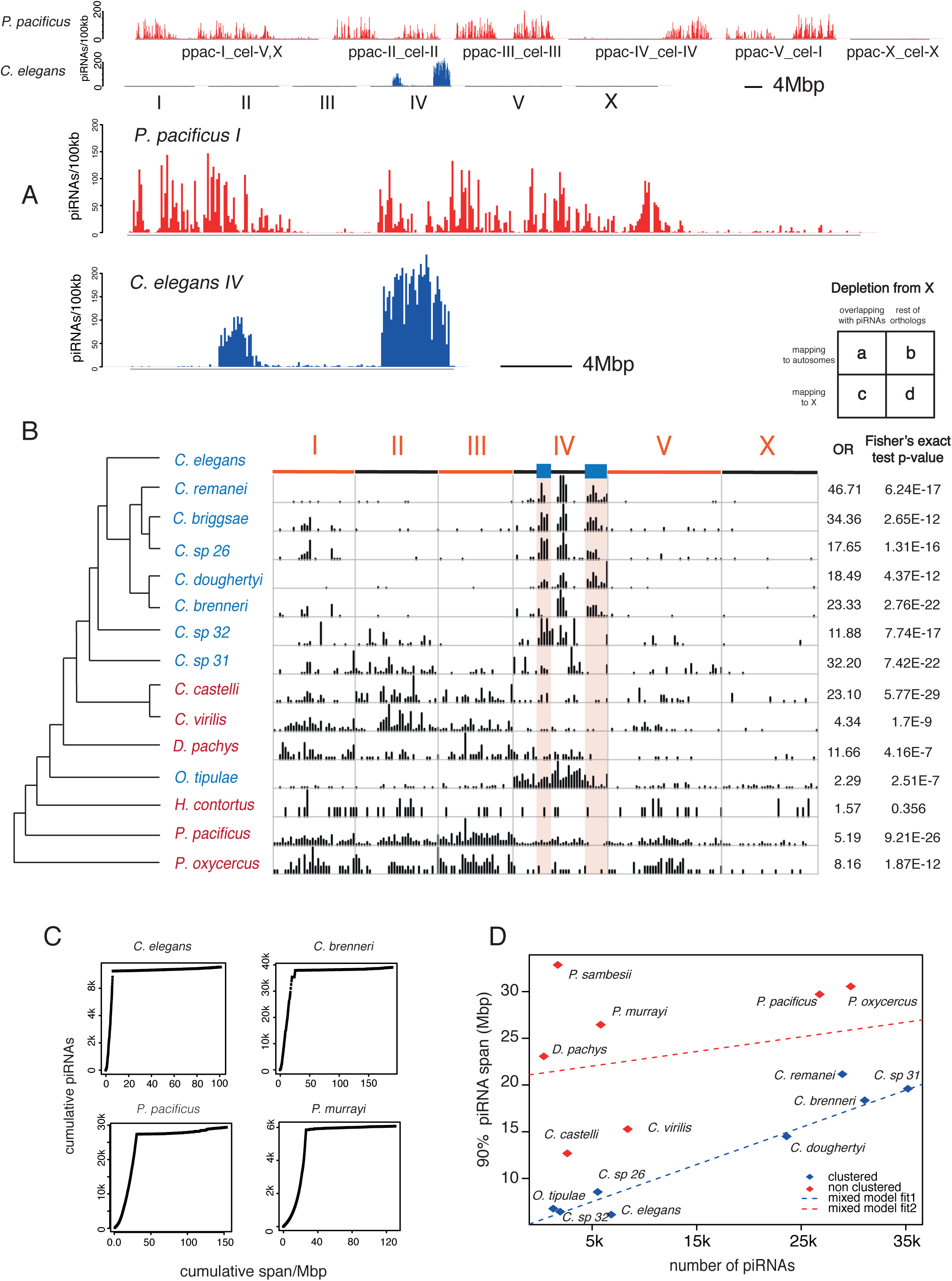
Two modes of piRNA locus organization in nematodes. A. Genomic distribution of piRNAs in *P. pacificus* (dispersed piRNAs) and *C. elegans* (clustered piRNAs). B. Mapping of piRNA regions in multiple nematode species to the *C. elegans* genome. Odds ratio (OR) and Fisher’s exact test p-value for X chromosome depletion is tabulated on the right hand side. C. Cumulative number of piRNAs against cumulative span of contigs ranked by the significance of piRNA enrichment. Examples of clustered species (*C. elegans, C. brenneri*) and non-clustered species (*P. pacificus, P. murrayi*). D. Discrimination plot of nematode genomes based on the number of piRNAs and the span of piRNA regions. Species were classified as clustered (blue dots) or non-clustered (red dots) on the basis of mapping of piRNA regions to *C. elegans* in B. Best-fit lines are from bimodal regression analysis using all points as input.

Most other nematode genome assemblies are of lower quality than those of *C. elegans* and *P. pacificus.* To analyse species with more fragmented genome assemblies we developed and validated a method to identify piRNA locus-dense contigs (Figure S3A,B). We then asked if it was likely that the clusters were in conserved regions across species, by using 1:1 orthologous protein coding genes to map the contigs to the *C. elegans* genome. We observed two large blocks of piRNA regions that contained orthologues that mapped to *C. elegans* Chromosome IV in the majority of *Caenorhabditis* species and also in *O. tipulae* (blue in Figure 3B). This pattern is consistent with the conservation of a small number of piRNA gene clusters in these species. In other nematodes, including *P. pacificus,* regions with elevated piRNA gene densities contain orthologues that map to several *C. elegans* chromosomes (red in Figure 2B). Across all nematodes, there was a depletion of piRNA loci linked to protein coding genes currently residing on the *C. elegans* X chromosome (Figure 3B). X linkage of sequence scaffolds was confirmed independently in three species: *O. tipulae,* where sex-linked genes have been used to identify the X chromosome (M.A-Felix, personal communication), in *H. contortus,* for which there is a recent chromosomal assembly (James Cotton, unpublished data), and in the chromosomal assembly of *P. pacificus* (Figure S3C).

We quantified piRNA clustering by examining the extent to which piRNA loci were concentrated within genomic regions (Figure 3C). Bimodal regression analysis identified two groups, one with more clustered piRNAs and one with less clustered piRNAs. These groups were congruent with those recovered from mapping to orthologous regions in *C. elegans* (Figure 3D). On the basis of these analyses we propose that there are two distinct modes of organisation of piRNA loci in nematodes, one clustered, similarly to *C. elegans* and one dispersed, similarly to *P. pacificus.*

### Clustered and dispersed piRNA loci have different chromatin environments

To further characterise the differences between clustered and non-clustered modes of piRNA gene organisation, we examined the chromatin environment of piRNA loci in *C. elegans* and *P. pacificus.* In the germline, the *C. elegans* genome is organised into mutually exclusive, stable domains of H3K27me3 repressive chromatin (“regulated domains”) and H3K36me3 transcriptionally active chromatin (“active domains”) (Evans et al., 2016). The majority (95%) of piRNA loci within the clusters overlapped with H3K27me3/regulated domains and only 5 of 9735 21U-RNA sequences were produced from H3K36me3/active domains (Figure 4A). Regulated domains were significantly larger within piRNA cluster regions compared to the rest of the genome (Figure S4A). piRNA loci were 227-fold depleted from active chromatin compared to a uniform distribution across piRNA cluster regions (empirical p<0.01). While chromatin domains have been defined based on analyses of the early embryo, they are present in the adult germline and show similar ChIP-seq profiles (Evans et al., 2016) (Figure S4B). Genes that overlapped with piRNA loci showed extremely low or null expression in the germline, while genes depleted of piRNA loci within cluster regions were highly expressed (Wilcox unpaired test p<2.2e-16, Figure S4C). Together these data strongly suggest that expression of *C. elegans* piRNAs occurs predominantly from regulated domains enriched in H3K27me3.

**Figure 4.**
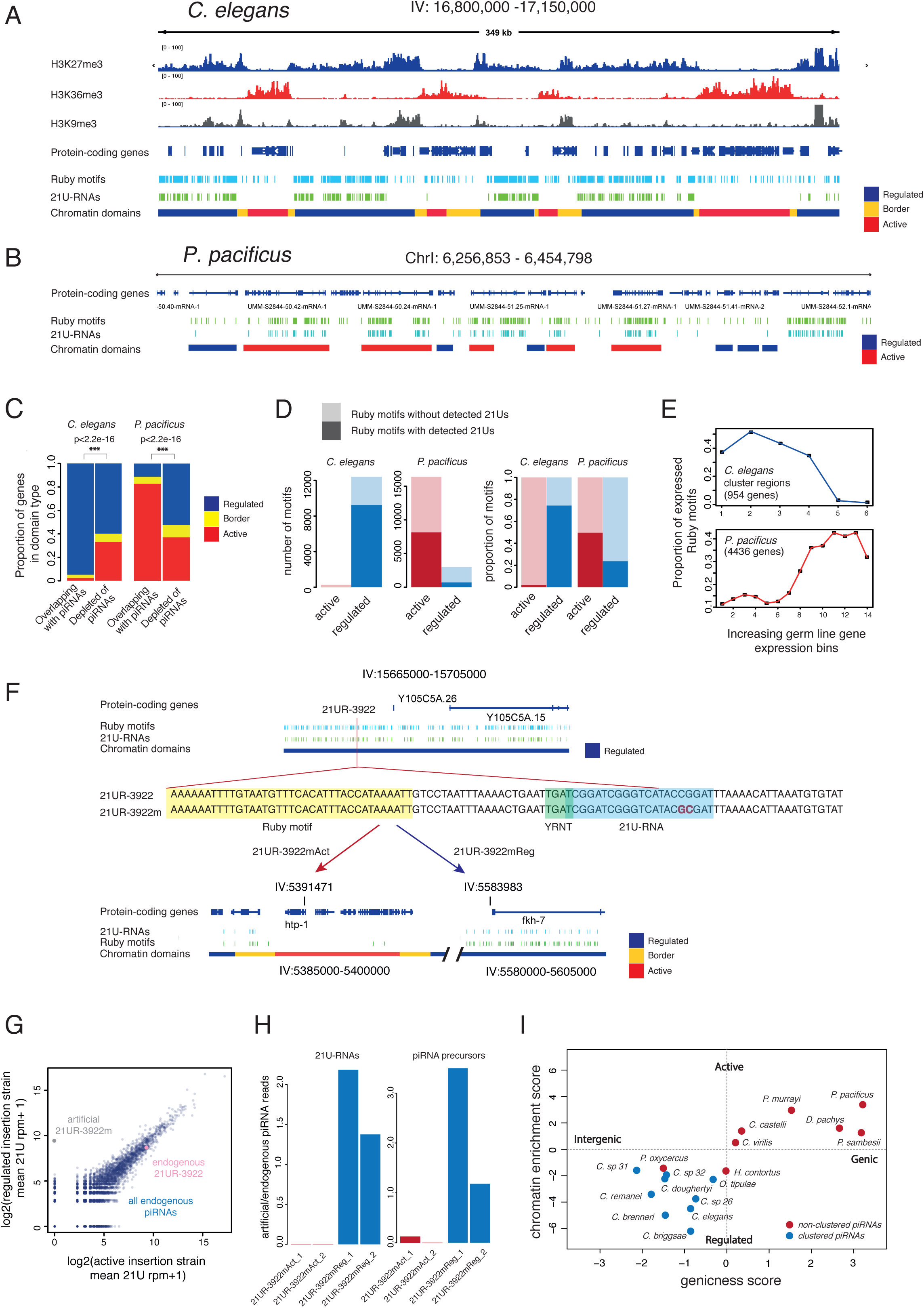
piRNAs in *C. elegans* and *P. pacificus* occupy different chromatin environments. A. *C. elegans* piRNA cluster regions are organised in multiple subclusters of piRNAs associated with H3K27me3 chromatin and interrupted by H3K36me3 chromatin domains depleted of piRNAs. B. *P. pacificus* piRNA loci are found within the introns of genes within H3K36me3 chromatin domains. C. Germ line expression of genes overlapping with piRNAs in *C. elegans* and *P. pacificus.* D. Correlation between gene expression and piRNA expression in *C. elegans* and *P. pacificus.* E. The fraction of Ruby motifs expressed in different chromatin environments in *C. elegans* and *P. pacificus.* F. Design and generation of artificial piRNA loci by genome-editing. G. Expression of an artificial piRNA placed in either regulated or active chromatin compared to the endogenous locus. H. Two fundamental modes of piRNA organisation in nematodes. Discrimination plot based on genic/intergenic proportions of piRNAs and mappings of 1:1 orthologs to *C. elegans* chromatin domains. Species are coloured according to the designation in Figure 3.

The organisation of piRNA loci in *P. pacificus* contrasts with that of *C. elegans.* 88% of piRNA loci were localised within protein-coding genes (1.6-fold enrichment within genes compared to intergenic regions, p<0.01 simulation test, Figure S4D). *C. elegans* piRNAs were biased toward intergenic regions (1.34-fold depletion from genes, p<0.01 simulation test, Figure S4D). *P. pacificus* piRNA loci are distributed approximately equally in the sense and antisense orientation of the genes in which they are found, but 97% of the intragenic piRNA loci are found within introns (empirical p<0.01 relative to random simulation) (Figure S4C). The chromatin environment of piRNA loci in *P. pacificus* is also distinct from *C. elegans.* The *C. elegans* 1:1 orthologues of the *P. pacificus* genes that contained piRNA loci were found almost exclusively in active H3K36me3 chromatin regions (Figure 4B,C) that are highly expressed in the germline (Figure S4C). While we do not have corresponding chromatin state data for *P. pacificus,* these data strongly suggest that its piRNAs are associated with active, rather than repressed, chromatin.

We examined the fraction of Ruby motifs associated with expressed 21U-RNAs in *C. elegans* and *P. pacificus* in different chromatin environments. In *C. elegans,* 74% of motifs in regulated chromatin were expressed, contrasting with less than 2% of motifs in active chromatin (Figure 4D, Figure S4E). In *P. pacificus,* piRNA motifs within genes whose *C. elegans* orthologues were within active domains were 2-fold more likely to produce piRNAs than those associated with *C. elegans* orthologues in regulated domains (Figure 4D, Figure S4E). The fraction of expressed motifs in *P. pacificus* was positively correlated with the expression of the overlapping protein-coding genes (Figure 4E), suggesting that host protein-coding gene expression promotes expression of the 21U-RNAs.

To test the involvement of chromatin structure in piRNA biogenesis directly, we used CRISPR-Cas9 genome editing in *C. elegans.* We selected an endogenous piRNA locus (21UR-3922) and modified the 21UR sequence to generate an artificial piRNA that could be distinguished from all endogenous piRNAs by sequencing (21UR-3922m; Figure 4F). We inserted the artificial piRNA gene into adjacent regulated and active chromatin domains within the piRNA cluster at the centre of Chromosome IV (Figure 4F). When inserted into regulated chromatin the artificial piRNA was expressed at the same level as the endogenous piRNA (Figure 4G,H). However, there was no detectable expression of the mature 21U-RNA when inserted into active chromatin (Figure 4G,H). Thus a repressive chromatin environment is necessary for piRNA production in *C. elegans.*

To generalise these observations we assessed the intergenic and genic proportions and the predicted chromatin environment of piRNA loci across nematodes. Species could be divided into two groups, one with more genic piRNAs associated with *C. elegans* orthologs enriched for active chromatin and one with more intergenic piRNAs associated with *C. elegans* orthologs enriched for regulated chromatin. These categories overlapped strongly with the pattern of clustering. Species with clustered piRNAs had H3K27me3/regulated-enriched piRNAs, whilst less clustered species had H3K36me3/active enriched piRNAs (Figure 4I).

We propose that there are two fundamental modes of piRNA organisation in nematodes. “Caenorhabditis”-type (C-type) piRNAs are organized into dense clusters within repressive chromatin. These sub-clusters are then grouped together into a large “super-cluster”, for example the regions on chromosome IV in *C. elegans.* In contrast, “Pristionchus”-type (P-type) piRNAs are more widely distributed across the genome where they are found within the introns of germline-expressed genes, enriched for H3K36me3.

### Chromatin domain organisation of piRNAs is under selection in nematodes

These different modes of piRNA organisation led us to ask how selection is acting on them. Using wild-isolate, genome-wide single nucleotide polymorphism (SNP) data from the *C. elegans* Natural Diversity Resource (Cook et al., 2017), *C. briggsae* (Thomas et al., 2015) and *P. pacificus* (Rödelsperger et al., 2014), we explored the predicted effects of SNPs on Ruby motifs by measuring their effect on match to a position weight matrix of all motifs. The allele frequency spectrum was markedly different between motifs within the piRNA locus clusters and those outside for both *C. elegans* and *C. briggsae* (Figure S5A,B). In *C. briggsae* and *C. elegans,* alleles with a low motif score were much less likely to be the major allele (present in >90% of strains) within the cluster than outside the cluster, implying that piRNAs are under stronger selection within the cluster than outside (p<1e-3 for both, Fisher’s Exact Test, Figure 5A,B). Similarly, both within and outside the cluster, selection was stronger on piRNAs within H3K27me3 domains than H3K36me3 domains (Figure 5C).

**Figure 5.**
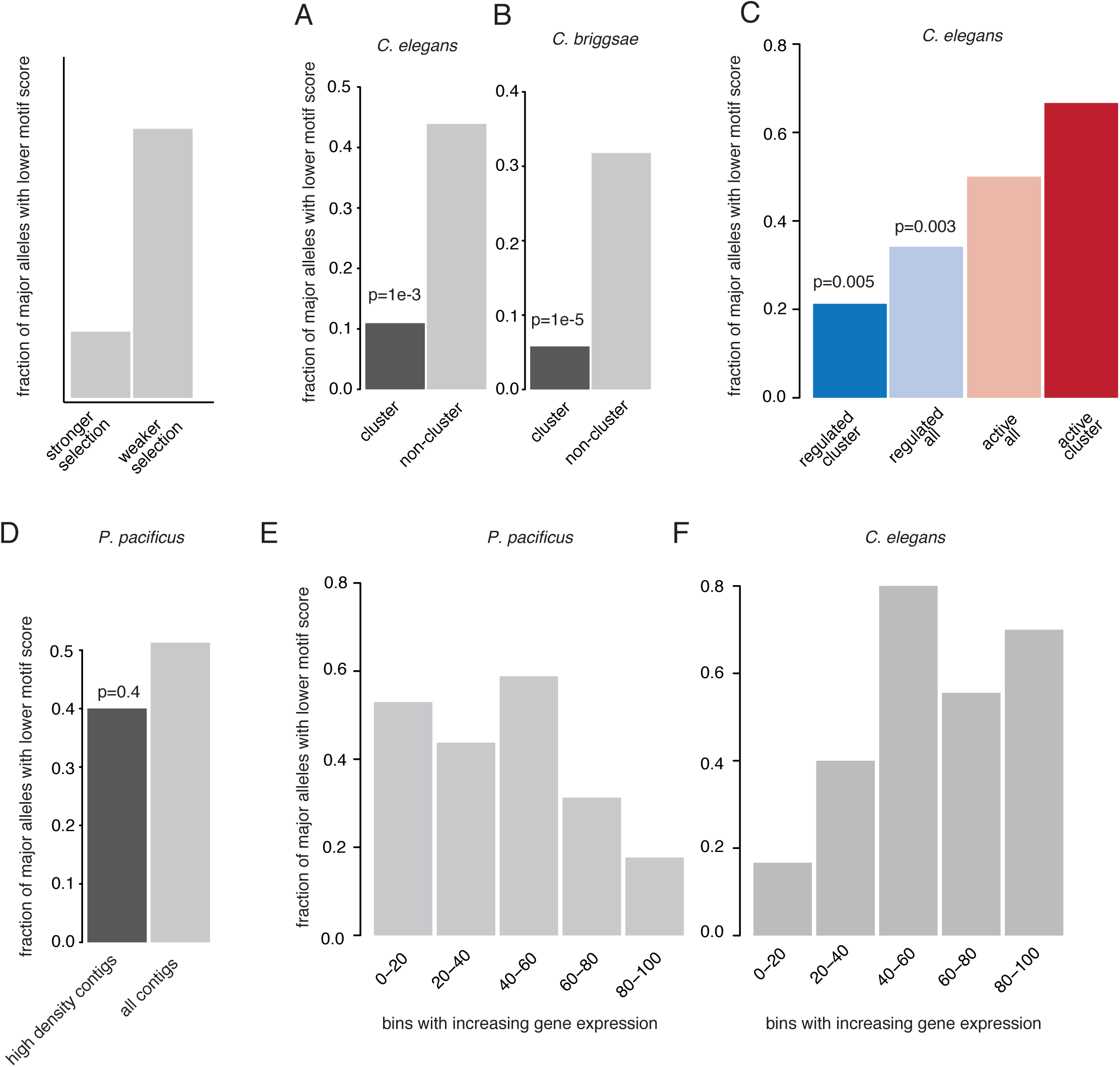
Selection at piRNAs is dependent on their chromatin environment. A-B. Fraction of high-scoring Ruby motifs that are the major allele (present in >90% of strains) in *C. elegans* (A) and *C. briggsae* (B). C.Frequency of the major allele of high-scoring Ruby motifs across different chromatin environments in *C. elegans.* D. Fraction of high-scoring Ruby motifs that are the major allele in regions of low and high piRNA density in *P. pacificus.* E. Fraction of Ruby motifs that are the major allele in genes stratified according to expression in *C. elegans* and *P. pacificus.*

In *P. pacificus,* observed SNPs tended to have a smaller effect on the motif score (Figure S5C), perhaps reflecting the lower information content within the motif (Figure 1). The major allele frequencies of low-scoring motifs did not vary with the density of piRNA loci, consistent with the lack of clusters (Figure 5D). However, in *P. pacificus,* SNPs predicted to disrupt piRNAs were less likely to be the major allele in highly expressed genes compared to lowly expressed genes (Figure 5E). Exactly the opposite trend was observed in *C. elegans* (Figure 5F). Together, these data confirm that the two different modes of piRNA biogenesis are under selection in their respective species.

### Comparative analysis implicates RNA Pol II pausing in piRNA production in nematodes

A puzzling feature of piRNA biogenesis in nematodes is how Pol II is regulated to produce the ∼30 nt precursor of mature piRNAs (Gu et al., 2012). Our finding that piRNAs can be found in both repressed and active chromatin environments is challenging, as *Pristionchus*-like piRNAs localise to regions of high transcriptional activity expected to favour elongating polymerase. Pol II is known to pause at the transcription start site (TSS) at certain types of eukaryotic protein-coding gene promoters. This results in the production of short-capped RNAs that are rapidly degraded by the nuclear exosome. This has been proposed as the underlying mechanism of piRNA transcription in nematodes (Gu et al., 2012) but so far evidence has been lacking.

Previous genome-wide analyses in *D. melanogaster* and human cell lines identified a sequence signature associated with RNA Pol II pausing characterised by a region with increased Tm relative to the genome-wide background and a region with reduced Tm immediately 3′ (Gressel et al., 2017; Nechaev et al., 2010). This architecture is thought to cause RNA Pol II to backtrack from the 3′ low T_m_ region to the 5′ high 0054_m_ region where RNA-DNA interactions are more stable. Consequently RNA Pol II pausing is observed at the boundary between the high T_m_ and low T_m_ regions. We examined T_m_ signatures around nematode piRNA TSSs. *P. pacificus* piRNA loci displayed a strong match to the pausing-associated sequence signature, such that a marked reduction in Tm relative to the genome-wide background coincided exactly with the predicted 3′ ends of piRNA precursors (Figure 6A,B; Figure S6A). Sequence motifs have not previously been identified downstream of *C. elegans* piRNAs (Ruby et al., 2006); nevertheless we found a clear match to the pausing-associated signature at *C. elegans* piRNA loci, although this was less pronounced than in *P. pacificus* (Figure 6A,B; S6A,B).

**Figure 6.**
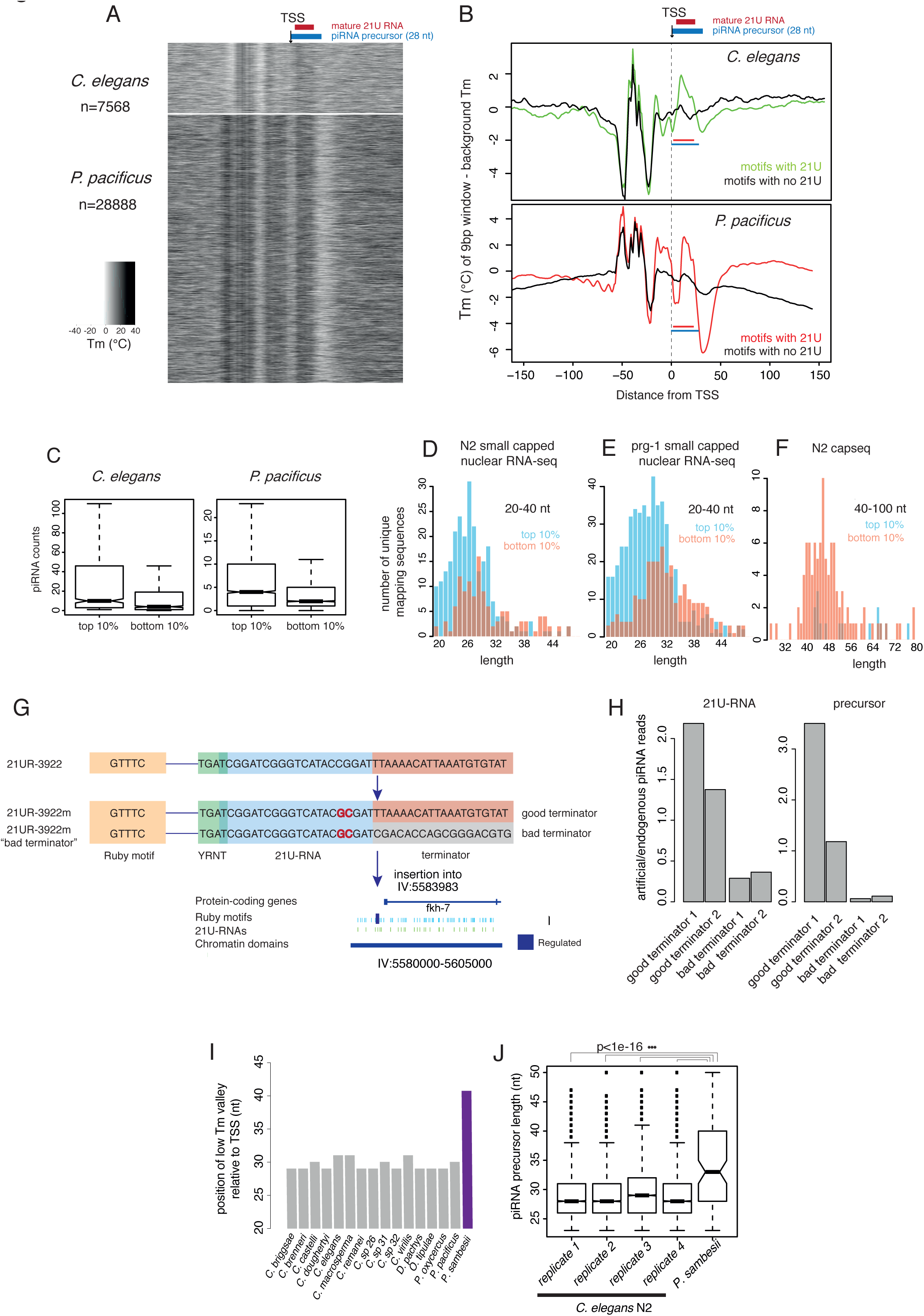
Transcription of short capped piRNA precursors in nematodes occurs through pausing of RNA polymerase II. A-B. Melting temperature profiles around piRNA transcription start sites in *C. elegans* and *P. pacificus.* C. Relationship between strength of pausing-associated sequence signature and mature piRNA levels. D-E. Length and abundance of piRNA precursors produced from loci with strong (top 10%) or weak (bottom 10%) pausing-associated signatures in *C. elegans* assessed by high-throughput sequencing of capped RNAs less than 50bp in size. D shows WT and E shows *prg-1* mutants. F. Longer nuclear Capped RNAs (40-60bp) emanating from piRNA loci with strong (top 10%) or weak (bottom 10%) pausing-associated sequence signatures assessed by high-throughput sequencing (compare with D and E). G. Design and generation of artificial piRNA loci with good and bad terminator sequences. H. Expression of the artificial piRNA loci with good and bad terminators. I. Location of the low melting temperature region downstream of piRNA loci. The distance to the centre of the low T_m_ “valley” from the piRNA transcription start site is shown. J. piRNA precursor length in *C. elegans* and *Plectus sambesii.*

We then examined the role of the pausing-associated sequence signature on piRNA expression by stratifying piRNA loci according to the strength of this signature. In *P. pacificus,* mature 21U-RNA expression was significantly greater from the loci with the strongest pausing-associated signatures than from the weakest (p<1e-10 Wilcoxon unpaired test) (Figure 6C). This effect was less marked but still significant in *C. elegans* (Figure 6C). The GC content of the 21U-RNA is not responsible for this effect, as sequence downstream of the 21U itself alone had a significant impact on mature piRNA abundance (Figure S6C).

To test the effect of the pausing signal on piRNA transcription, we examined piRNA precursors from nuclear short capped RNA sequencing data from a previous study (Weick et al., 2014). piRNA precursors produced from loci with strong pausing signals were significantly shorter and on average two-fold more abundant than those produced from loci with weak pausing signals (Figure 6D). This effect was emphasized in *prg-1* mutants, where mature piRNAs are not present, leading to accumulation of piRNA precursors in sequencing data (Figure 6E, S6D-F). Consistently, the few atypical longer precursors (40-100bp) sequenced by capseq (Gu et. al., 2012) mapped almost exclusively to loci with the weakest pausing signals (Figure 6F, S6D-F).

We modified the sequence downstream of the artificial 21UR-3922m to increase the Tm by 27.5°C, and inserted this new artificial piRNA (“21UR-3922m-bad terminator”) into the same site within the H3K27me3 domain (Figure 6G). This allowed us to compare of the expression of two piRNA loci that have different pausing-associated sequence signatures but are otherwise identical. The 21U-RNA with the higher downstream Tm was expressed at around 3-fold lower levels than the endogenous piRNA with the same sequence (Figure 6H). Moreover, the piRNA precursor was expressed at ∼10-fold lower levels than the endogenous piRNA precursor (Figure 6H). Thus the Tm of the downstream sequence alone is sufficient to affect transcription of the piRNA locus.

We detected similar Pol II pausing-associated sequence signatures across nematodes (Figure 6I, Figure S7-4). In *P. sambesii,* the low T_m_ region downstream of the 21Us is shifted ∼10 nt further downstream compared to the rest of nematodes examined (Figure 6I, Figure S7-4). piRNA precursors are significantly longer in this species compared to *C. elegans* (Figure 6J), consistent with a role for the pausing signal in piRNA transcription termination.

To test the involvement of pausing in piRNA production we examined *C. elegans* mutants carrying a deletion in TFIIS. TFIIS is a general transcription elongation factor that rescues backtracked Pol II complexes by promoting cleavage of the 3′ end of the RNA by Pol II itself (Schweikhard et al., 2014). In *D. melanogaster,* TFIIS deletion leads to an increase in the length of promoter-associated short capped RNAs, indicative of pausing and backtracking of Pol II at the promoter (Nechaev et al., 2010). *C. elegans* mutants carrying a deletion in the orthologue of TFIIS (*T24H10.1* ok2479) showed significantly longer piRNA precursors (Figure 7A, S7-1 A-D) and a modest, but consistent, decrease in mature piRNA abundance (Figure 7B). The effect of the TFIIS deletion is specific for short-capped RNAs, as we did not find differences in length distributions of rRNA degradation fragments (Figure S7-2 A-D) or tailed 22G-RNAs (Figure S7-2 E). To test whether TFIIS promotes the cleavage of piRNA precursors, we examined sequencing data for potential 3′ cleavage fragments, with a 5′ monophosphate mapping 28-38 nucleotides downstream of the piRNA transcription start site. We detected potential cleavage products from around 10% of loci in wild type nematodes, but these products were 2-fold reduced in TFIIS mutants (Figure 7 C,D). These data support a direct role for Pol II pausing in proper termination of piRNA transcription and release of piRNA precursors.

**Figure 7.**
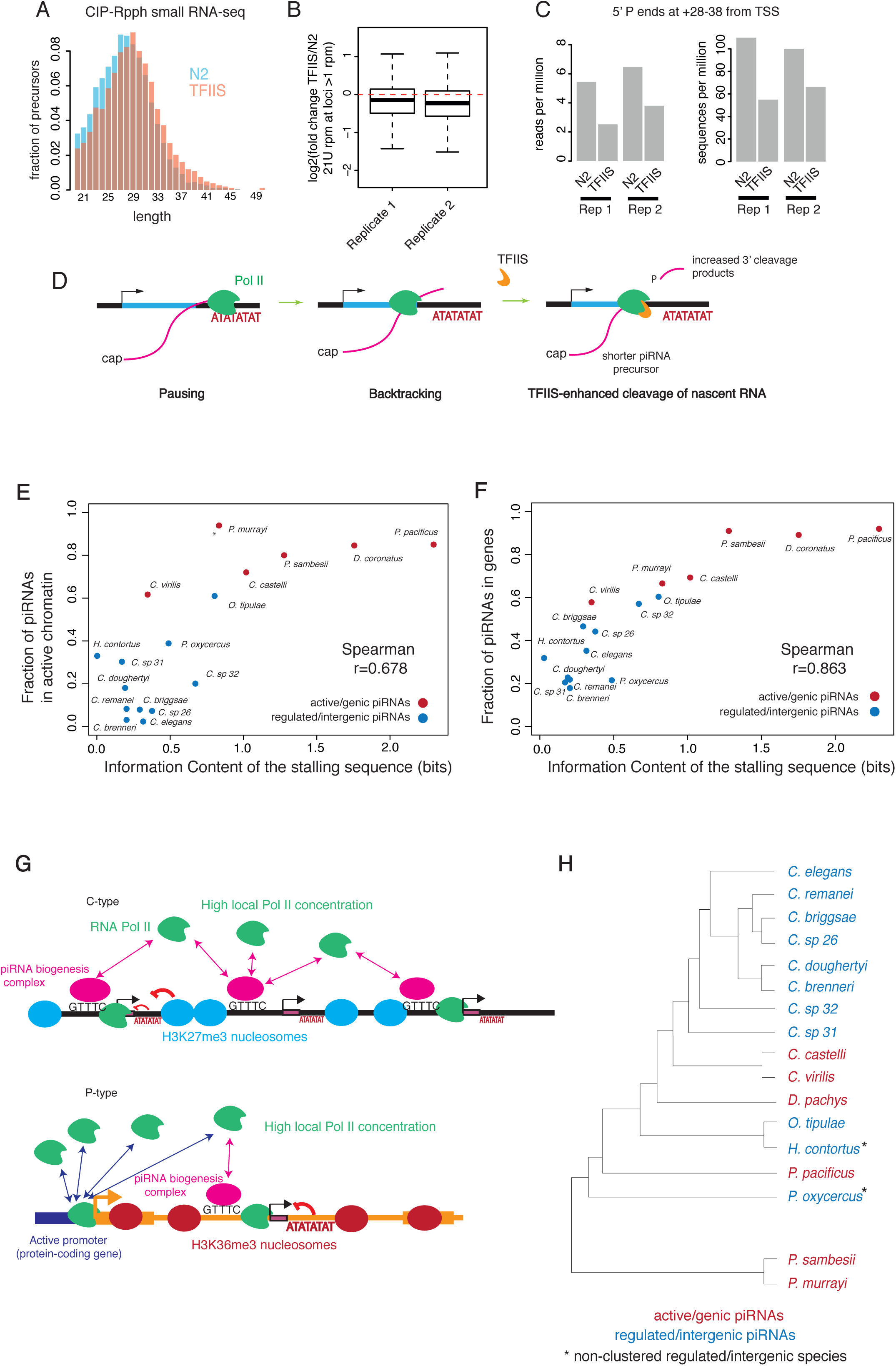
Chromatin and RNA Pol II pausing together control nematode piRNA biogenesis. A. piRNA precursor length in TFIIS mutants compared to wild type. B. Mature piRNAs in TFIIS mutants compared to wild type. C. 3′ degradation fragments of piRNA precursors in TFIIS compared to WT *C. elegans.* 3′ degradation fragments are defined as 5′ monophosphate small RNAs whose 5′ end maps between 28 and 38 downstream of the piRNA transcription start site. D. Model for how Pol II pausing contributes to piRNA biogenesis. The AT-rich region downstream of the 21U sequence causes Pol II to destabilise and stall, leading to backtracking and TFIIS-mediated cleavage of the 3′ ends of nascent RNAs. This leads to a decrease in piRNA precursor length and an increase in the abundance of 3′ degradation fragments, and influences 21U levels by altering Pol II dynamics at piRNA promoters.E-F. Relationship between the average strength of pausing signal and enrichment of piRNAs in active chromatin (A) and enrichment of piRNAs in genic regions (B) across different nematode species. G. Model for piRNA transcription in C-type and P-type nematodes. H. Widespread phylogenetic distribution of active-genic and regulated-intergenic piRNAs.

In *C. elegans* piRNA precursors are trimmed by the exonuclease PARN-1, an orthologue of Trimmer from *B. mori* (Tang et al., 2016). An alternative explanation for our findings would be if the sequence signature exerted its effect post-transcriptionally by affecting the trimming of capped piRNA precursors. However, this is unlikely to be the case as capped piRNA precursor length differences between loci with strong and weak pausing signals remain unaffected in the absence of trimming (Figure S7-3B). In addition, the overall length of capped piRNA precursors does not change in *parn-1* (tm869) mutants compared to wild-type nematodes (Figure S7-3C). Finally, the increase in precursor length due to TFIIS deletion is still significant when assessed in a *parn-1* (tm869) background (Figure S7-3D). This data is consistent with cytoplasmic trimming on an uncapped Piwi-bound piRNA precursor, which has been shown in *B. mori* (Izumi et al., 2016). Therefore, nuclear short capped RNAs represent early unprocessed piRNA precursor transcripts.

Altogether, our data strongly suggests that the sequence properties of piRNA loci promote RNA Pol II pausing and are thus important for generation of short piRNA precursors.

### Synergy between chromatin environment and pausing-associated sequence signals in nematode piRNA biogenesis

Though clearly important for piRNA biogenesis, the Pol II pausing signature is weaker in *C. elegans* than in *P. pacificus.* To test whether this extended to the remaining C-type and P-type species, we quantified the strength of Pol II pausing-associated sequence signatures at piRNA loci across nematodes (Figure S7-4). Across nematode species the strength of the Pol II pausing-associated sequence signature correlated positively both with the proportion of piRNAs in active chromatin and the proportion of piRNAs within genes (Spearman rho > 0.67 for both, p=1.9e-3 and p=3.89e-6; Figure 8E,F). Thus C-type species, in which piRNA loci are found within H3K27me3 chromatin domains, have relatively weak pausing signals while stronger pausing signals are found in P-type species where piRNA loci are found within H3K36me3 domains.

## Discussion

Our analysis of piRNA organisation and biogenesis across nematodes not only illuminates the evolutionary history of the piRNA system in nematodes but also provides unexpected insights into the fundamental mechanism of piRNA production in *C. elegans.*

### piRNA transcription evolved from snRNA transcription

The Ruby motif upstream of *C. elegans* piRNAs was derived from the 3′ half of the ancestral nematode SNAPc motif upstream of snRNA loci. In *C. elegans,* SNAPc binding sites are globally enriched in piRNA clusters but do not seem to localise at a particular distance from piRNA TSSs (Kasper et al., 2014). Our analysis predicts that SNAPc binds directly upstream of piRNA loci to drive Pol II-dependent transcription of piRNAs in *P. sambesii* and *P. murrayi.* Even in ancestral nematodes that have lost the piRNA pathway, the SNAPc motif upstream of snRNA loci closely resembles the *P. sambesii* piRNA motif. Thus nematode piRNA biogenesis evolved from snRNA transcription. The separation of Ruby and SNAPc motifs in rhabditid nematodes (Clades III, IV, V) may correlate to the appearance of specialised piRNA factors such as PRDE-1, which are not conserved in plectids. PRDE-1 interacts with SNPC4 (Kasper et al., 2014) and may recruit SNAPc to the *C. elegans* Ruby motif in the absence of direct DNA binding.

### RNA Pol II pausing as a key feature of piRNA production in nematodes

Previous identification of piRNA precursors by deep sequencing in *C. elegans* demonstrated piRNA production by RNA polymerase II (Billi et al., 2013; Gu et al., 2012), and it was hypothesized that piRNA precursors might result from promoter-proximal Pol II pausing (Gu et al., 2012). This suggestion was prescient: our analysis of piRNA biogenesis across nematodes reveals that this process is indeed likely to be responsible for the short length of piRNA precursors.

Following transcription initiation, Pol II forms an early elongation complex with limited elongation capacity, requiring further regulation to transition into productive elongation. During early elongation, Pol II is highly susceptible to pausing, which can lead to backtracking by a few nucleotides (Schweikhard et al., 2014). Release of the stalled complex requires cleavage of the 3′ end of the transcript, promoted by TFIIS. At many promoters Pol II rarely switches into elongation and is instead released from the template. This process results in accumulation of a short, capped RNAs derived from the promoter of the gene. We suggest that exactly this cycle of backtracking, cleavage and release of RNA results in the production of piRNA precursors. Our model is supported by recent data showing that binding and release of Pol II at transcription start sites and the production of short capped RNA occurs continuously at paused loci (Krebs et al., 2017).

The involvement of Pol II pausing in production of piRNAs in *C. elegans* is consistent with other findings demonstrating production of functional small RNAs from promoter-proximal Pol II. Mammalian Ago-2 binds to small RNAs produced from hairpins derived from paused Pol II at protein-coding gene promoters (Zamudio et al., 2014). In ciliates, Pol II pausing in conjunction with TFIIS activity generates ∼25 nt RNAs that are loaded into Piwi proteins (Gruchota et al., 2017). These observations suggest that Pol II pervasive initiation and pausing can be co-opted for the biogenesis of functional small RNAs across eukaryotic genomes, including mammals.

### Dynamic evolution of the piRNA pathway: two alternative modes of piRNA organisation and biogenesis across nematodes

Here we show that two modes of piRNA biogenesis are found in nematodes (Figure 8G,H). in C-type species piRNAs are found clustered together at high density, within regions of H3K27me3-rich chromatin, as in *C. elegans. C. elegans* piRNA clusters are therefore better described as “clusters-of-clusters” within H3K27me3 domains separated by H3K36me3 rich regions depleted of piRNAs. Contrastingly, in P-type species piRNAs are found in active chromatin, and are not clustered together, as observed in *P. pacificus.* The piRNA types also differ in their reliance on cis-acting Pol II pausing-associated sequence signatures downstream of the piRNAs: whilst these signatures are relatively weak at piRNA loci in C-type species, they are much more obvious at piRNA loci in P-type species.

piRNA precursors must be transcribed at a sufficient rate whilst ensuring that Pol II does not switch to elongation mode. We suggest that the two modes of piRNA biogenesis in nematodes represent alternative solutions to this problem. The high density of piRNAs in C-type species may serve to attract sufficient Pol II to a region devoid of active genes (Figure 7B). Once present in the H3K27me3-rich chromatin surrounding piRNA loci, Pol II is unlikely to switch to elongation. A contrasting situation occurs in P-type species. Here, we suggest that piRNA promoters “piggyback” on the capacity of protein-coding genes to attract Pol II, and thus a high piRNA density is not required. However, Pol II is prone to switch to elongation within these regions so a stronger pausing-associated sequence is required downstream of the piRNA to ensure correct termination.

This model is consistent with genome-wide studies in a variety of model systems showing that H3K27me3 is anticorrelated with the elongating form of Pol II. It has been reported that H3K27me3 accumulation at promoters in the absence of H2K119 mono-ubiquitination by PRC1, directly promotes short capped RNA production at poised genes (Min et al., 2011). This is particularly interesting because *C. elegans* lacks a clear germline-expressed ortholog of PRC1 and thus likely do not have germline H2B-monoubiquitination in H3K27me3 rich regions (Karakuzu et al., 2009). However, this finding applies to bivalent genes enriched in H3K4me3 in addition to H3K27me3 and whether H3K27me3 alone is sufficient is unknown.

Currently we have insufficient evidence to establish which mode of biogenesis was ancestral. It appears that the mode may switch dynamically. For example *D. pachys* has P-type piRNAs despite being nested within the C-type species *C. elegans* and *O. tipulae.* It is therefore possible that both types of piRNAs use common biogenesis machinery and could coexist within the same organism. In support of this we note that some species are “intermediate” between clustered and non-clustered organisation (Figure 3F), and there is a gradation of both the percentage of piRNA loci in genes and the strength of the pausing-associated sequence signature downstream of the piRNAs (Figure 7E,F).

Overall, the novel aspects of piRNA biogenesis that we describe for *C. elegans* illustrate the utility of using a cross-species approach to investigate mechanistic questions in model organisms. We predict that this approach will be similarly important in addressing fundamental questions of mechanisms for other pathways of epigenetic regulation and genome defence, in part because these systems evolve rapidly across species.

## Methods

### Nematode culture and RNA extraction

*C.elegans, Oscheius tipulae* and *Diploscapter pachys* were grown at 20°C in standard nematode growth medium (NGM) agar plates feeding on HB101 *E. coli* seeded in LB medium. *P. oxycercus* was grown in the same way at 25°C. *P. sambesii* was grown at 25°C in low salt NGM feeding on HB101 seeded in water. *H. bacteriophora* strain M31e was grown on lipid agar media using *Photorhabdus temperata* TRN16 as a food source. To enrich for germline tissue, *C. elegans, O. tipulae* and *H. bacteriophora* nematodes were synchronised by hypochlorite treatment and grown to adulthood. *P. sambesii* adults were selected using the COPAS sorter (Union Biometrica). Nematodes were resuspended in Trizol reagent (Life Technologies) and tissues were disrupted by bead-beating except H. bacteriophora where tissues were disrupted by grinding. RNA was then extracted and precipitated from Trizol according to the manufacturer’s instructions.

### Genome sequence and assembly of P. oxycercus

Genomic DNA of *P. oxycercus* was prepared using a DNeasy Blood and Tissue kit (Qiagen), according to the manufacturer’s instructions. Library preparation and sequencing of P. oxycercus was performed by GATC Biotech.

Quality control of *P. oxycercus* raw genomic data was assessed using Fastqc v0.10.1, and reads were quality trimmed using skewer v0.2.2 (Jiang et al., 2014) with parameters −q 30 −| 51. K-mer plots were generated with kmc, which revealed extensive heterozygosity. A preliminary single-end assembly was generated with Velvet (Zerbino and Birney, 2008) and contaminants were identified using blobtools (Kumar et al., 2013). The data were digitally normalised to 80x coverage to facilitate assembly. Assembly was carried out with SPAdes v3.5.1(Bankevich et al., 2012) correcting the reads with BayesHammer within the SPAdes pipeline. Assembly parameters were −k 21,33,55,77,99 −cov-cutoff auto --careful. *Plectus sambesii* was assembled as described in Rosic. et. al., 2017.

Examination of the resulting distribution of contig read coverages revealed a bimodal distribution in both genomes suggesting heterozygosity. Haploid coverage contigs were collapsed and postprocessed with Redundans (Pryszcz and Gabaldón, 2016), that runs SSPACE3 (Boetzer et al., 2011) and SOAPdenovo Gapcloser internally. The identity percentage identity for the collapse of redundant contigs was chosen to be 90%, since lower values did not result in a significantly greater reduction of the span, and no decrease in the number of KOGs by CEGMA (Parra et al., 2007) was found. RepeatModeler v1.0.8 (Smit and Hubley *RepeatModeller open 1.0*) was used to generate species-specific repeat libraries that were concatenated with a nematode repeat library from RepBase. The combined library was used to mask the genome with RepeatMasker v4.0.7 (Smit, Hubley and Green *RepeatMasker open 4.0).*

To annotate the *P. oxycerca* genome, we generated rRNA-depleted paired-end total RNA-seq libraries. Quality control of the RNA-seq reads was performed with Fastqc and Skewer with parameters −q 30 −l 50. Reads were aligned to the assemblies with STAR v2.5.2 (Dobin et al., 2013) with default parameters except for --twopassMode Basic. Alignment bam files were used for automated annotation using BRAKER v1.9(Hoff et al., 2016).

### Small RNA library preparation and sequencing

RNAs were pretreated to modify 5′ ends in two ways. For RPPH treatment, 1-2 *μ*g of total RNA was treated with 10 units (2 *μ*l) RNA 5′ pyrophosphohydrolase (NEB) for 1 h at 37°C. The treated RNA was phenol-chloroform extracted and ethanol precipitated with sodium acetate and glycogen for 2 days, and resuspended in RNase-free water. For CIP-RPPH treatment, 3-5 *μ*g of total RNA was treated with 4 *μ*l of QuickCIP (Quick dephosphorylation kit, NEB) in a total volume of 40 *μ*l for 90-120 min at 37°C. RNA was phenol-chloroform extracted, precipitated overnight with sodium acetate and glycogen and resuspended in RNase-free water.

Small RNA libraries from treated or untreated RNA were built using the TruSeq small RNA kit from Illumina according to the manufacturer’s instructions except for an increase in the number of PCR cycles from 11 to 15. Libraries were eluted in 0.3 M NaCl, ethanol precipitated and quantitated with Qubit and TapeStation. Libraries were pooled in groups of 6 to 12 per lane and sequenced on an Illumina HiSeq2000. The Illumina universal adapter was trimmed from small RNA reads using cutadapt v1.10 and reads were mapped to the corresponding genome assemblies with Bowtie v0.12 (Langmead et al., 2009) with parameters −v 0 −m 1.

### Phylogenetic profiling of piRNA pathway genes

Identification of Piwi from nematode genomes was performed essentially as described (Sarkies et al., 2015). Briefly, reciprocal blastp searches were performed on predicted protein sets using *C. elegans* PRG-1 as a query sequence. In species where Piwi is not present, the best hit for PRG-1 is another Argonaute subfamily member. In species where we did not identify Piwi from predicted proteins (such as *C. plicata),* we additionally performed tblastn searches against the genome sequence using *C. elegans* Piwi as a query sequence, to verify the absence of the gene.

### De novo motif discovery and piRNA annotation

Sequences 110 bp upstream and 30 bp downstream of 21U mapping sites were extracted and subsets of 2000 sequences were randomly selected to predict motifs de novo using MEME v4.10.1 (Bailey et al., 2009) with parameters −dna −mod zoops −maxsize 2000000 -nmotifs 10. Genome-wide nucleotide content was used as a background model for MEME. The resulting motifs from each subset were compared to assess the consistency of the analyses, and the full dataset of upstream sequences was scanned with the obtained motif position weight matrix (PWM) using a custom Python implementation (scripts available upon request). For each species, the distribution of mapping positions showed a peak at the expected distance of 42 b from the 21U mapping site. The distributions of PWM scores showed bimodal distributions, indicating good separation of true piRNA motifs. These distributions were used to define species-specific threshold to select true positive motifs, which were annotated as piRNA loci. Motifs were used for genome-wide scans without 21U sequence information using the same threshold. In the case of *P. sambesii,* the increased length of the motif resulted in an exceptional specificity of genome-wide scans, so all motif positions were considered “true” piRNA loci. *P. murrayi* piRNAs were defined in the same way using the *P. sambesii* motif. In *H. bacteriophora* upstream sequences displayed no positional bias for GTTTC nor any of the motifs predicted by MEME. We performed motif discovery on other small RNA lengths and 5′ nucleotides as controls, finding no enrichment of motifs. To map early precursor transcripts, we mapped RPPH treated small RNA libraries to piRNA loci identifying reads containing 5′ and 3′ extensions to the 21U-RNA sequence as described previously (Weick et al., 2014).

### Annotation of snRNA genes and prediction of SNAPc motifs

We annotated SNAPc-dependent non-coding RNA loci across nematode genomes using Infernal v1.1.2 (Nawrocki and Eddy, 2013). RFAM alignments for U1, U2, U3, U4, U4atac, U5, U7, U11 and U12 snRNAs (Pol II-dependent), and U6 snRNA, U6atac snRNA, 7SK RNA and MRP RNA (Pol III-dependent) (see Supplementary tables for RFAM IDs) were used to build and calibrate Infernal covariance models with default parameters. Nematode genomes were searched with the trained models with default parameters. The 100 bp upstream of significant hits (E-value >1e-3) were extracted for motif prediction with MEME v4.10.2 (Bailey et al., 2009) with parameters −dna −mod zoops −nmotifs 10.

### piRNA clustering analysis and comparison to the C. elegans genome

We identified piRNA-enriched contigs by applying a binomial test against a null hypothesis of even distribution of piRNAs across the genome. Contigs were ranked based on the p-value of the enrichment, and a cumulative plot of piRNA counts by span was generated. This approach yielded a good separation between piRNA-enriched and piRNA-depleted contigs and allowed us to define the span covered by piRNAs for each species. We validated this method using a fragmented *de novo* assembly of the *C. elegans* genome, where we successfully recovered the genomic regions corresponding to the *C. elegans* piRNA clusters. We applied the same method to the genome of *O. tipulae,* for which there is a good sequence-anchored linkage map. The top 10 most enriched contigs were found in two blocks mapping to chromosomes OtIV and OtV, consistent with the presence of clusters similar to *C. elegans.* To quantify the extent of clustering we calculated the significance of the enrichment of each contig compared to the null hypothesis of a uniform distribution of piRNAs across the genome using a chi-squared test and ordered by decreasing significance of enrichment. We then calculated the cumulative frequency of total loci and identified the span required to reach 95% of the total loci. We then plotted this value against the total number of piRNAs to give the plot in Figure 3. Multimodal regression analysis was performed using the Mixtools package in R.

For each species, we defined gene orthology to *C. elegans* with Orthofinder v0.6.1(Emms and Kelly, 2015) running mcl 14.137. 1:1 orthologs were filtered from the final dataset (Supplementary tables, *all species-piRNA organisation),* and the *C. elegans* orthologs of genes localised in piRNA contigs for each species were identified.

### Analysis of the genomic organisation of piRNAs

We used Integrative Genome Viewer to visualise *C. elegans* piRNA positions relative to genes and a variety of chromatin tracks from the modENCODE project. The modENCODE accession numbers for H3K27me3 and H3K36me3 in early embryos and germline-containing adults are 6254, 6256, 5163 and 5165. The overlap of *C. elegans* piRNAs with *C.elegans* chromatin domains in early embryos and L3 larvae (Evans et al., 2016) was calculated with Bedtools v2.25.0 (Quinlan and Hall, 2010). To estimate chromatin environment of piRNA loci in other species, 1:1 orthologs were selected as described above, and annotated for chromatin state based on the state of the *C. elegans* ortholog. We tested the association of piRNA-containing genes with particular chromatin domain types with a Fisher’s exact test. We defined the “chromatin enrichment score” as the log2 of the odds ratio of a 2×2 table of genes with and without piRNA locus association in active versus regulated chromatin. The germ line expression of these groups of genes in *C. elegans* was assessed using RNA-seq from dissected germ line tissue (Ortiz et al., 2014), and differences in expression were tested with a Wilcoxon rank sum test.

The localisation of piRNAs relative to genes and intronic regions in each genome was calculated using bedtools intersect v2.25.0. We randomly simulated piRNA positions genome-wide to generate null distributions, and simulated piRNA positions within their contig of origin to account for potential regional biases in those genomes We defined a “genicness score” for each species as the genic/intergenic ratio of piRNA locus positioning in the real dataset divided by the median of the 100 genic/intergenic ratios of simulated datasets.

### Expression of artificial piRNA loci

We used CRISPR-Cas9 to insert an artificial piRNA locus with a 21U-RNA sequence absent from the *C. elegans* genome. We based the artificial piRNA on the endogenous piRNA 21UR-3922 (IV:15671231-15671251) and edited the mature piRNA sequence such that it had no match to the *C. elegans* genome (TCGGATCGGGTCATACCGGAT > TCGGATCGGGTCATACGCGAT), leaving the upstream and downstream parts the same (mature 21-RNA sequence in capital letters):

aaaaaattttgtaatgtttcacatttaccataaaattgtcctaatttaaaactgaattgaTCGGATCGGGTCATACGCGATttaaaacattaaatgtgtat

This construct was inserted into a regulated domain (IV:5583983), or into the first intron of *htp-1* within active chromatin (IV:5391471) by injecting preformed Cas9-gRNA complex along with the appropriate repair oligo (Table 1), with *dpy-10* as a coinjection marker (Paix et al., 2015). Correct insertions were verified by PCR (Table 1) and Sanger sequencing.

We modified the region immediately downstream of the mature piRNA sequence to increase its GC content, and thus the melting temperature of the region, and inserted this new locus (21UR-3922m “bad terminator” into the same location within the regulated domain (IV:5583983).

Two strains carrying each insertion representing independent insertion events were generated. The strains were synchronised by hypochlorite treatment and grown from L1 for 72 h for RNA isolation, and RPPH libraries were prepared as described above. We recovered 21U-RNA and piRNA precursor counts for each piRNA locus. We used DEseq2 to estimate library size factors and identify differentially expressed 21U-RNAs. Additionally, we normalised the mature piRNA and precursor read counts of the artificial locus relative to those of the endogenous locus.

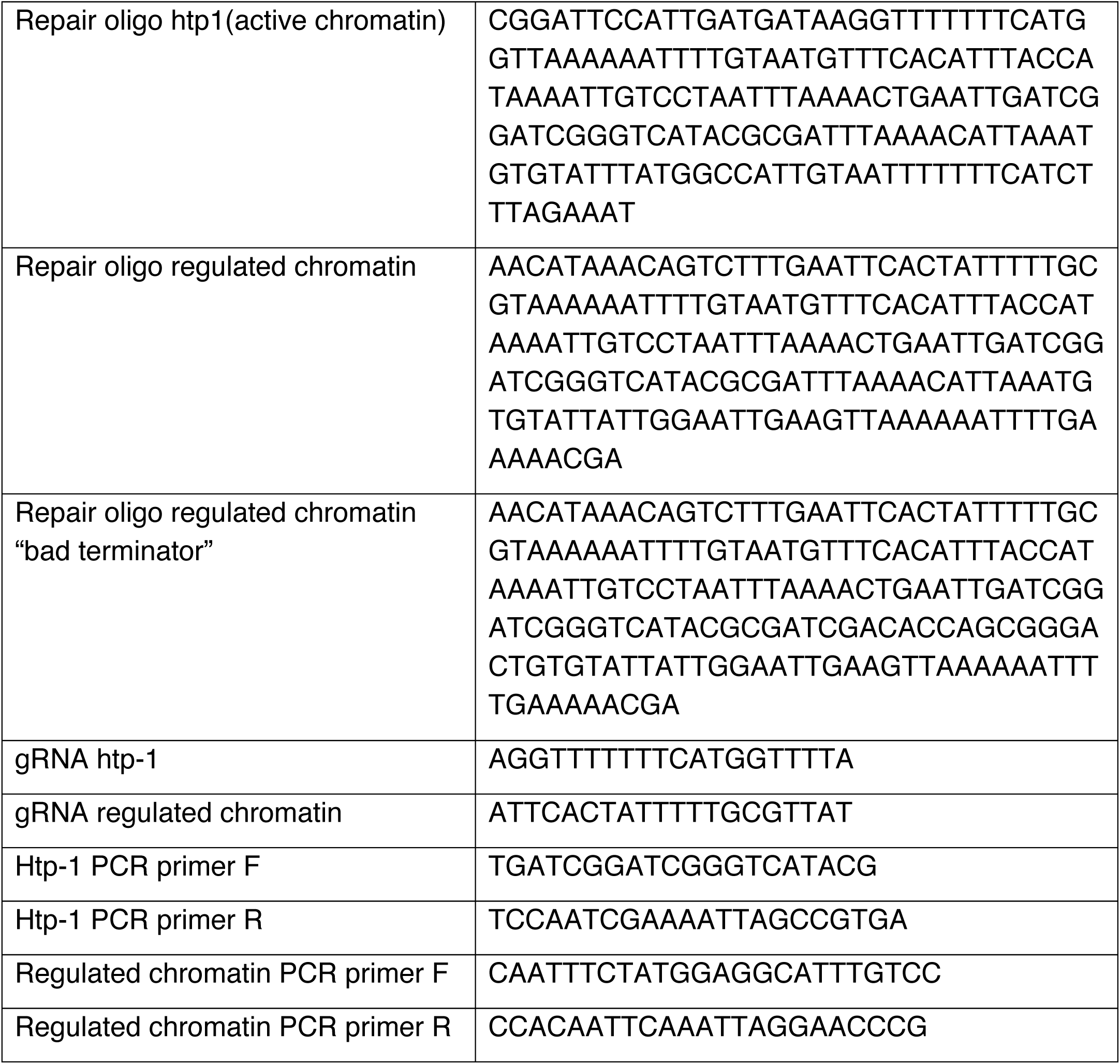

### Effect of SNPs on piRNA motif scores

We downloaded genomic variation data for *C. elegans* from the *C. elegans* natural diversity resource (CeNDR, https://www.elegansvariation.org/,(Cook et al., 2017). *C. briggsae* SNPs were taken from (Thomas et al., 2015). *P. pacificus* SNPs were downloaded from the *Pristionchus* variome site (http://www.pristionchus.org/variome/, Rödelsperger et al., 2014). To establish the impact of SNPs on piRNA motifs, we identified Ruby motifs using the specific position weight matrices (PWM) and selected motifs with a score >=500 log odds ratio and recalculated the motif scores after substituting any SNPs. For piRNA loci with a log odds ratio >1000, we identified SNPs with a predicted effect of >=400 change in log odds score (C. *elegans, C. briggsae*) or >=200 for *P. pacificus* (due to lower motif information content). We determined whether SNPs affecting piRNAs were major alleles (present in >90% of strains) or minor alleles (present in <10% of strains). We explored their association with annotated piRNA clusters, chromatin domains and genes using Bedtools for data integration.

### T_m_ analysis of piRNA loci and their effects in piRNA transcription

Sequences 200 bp upstream and downstream of piRNA motifs were extracted using the Bedtools module getfasta v2.25.0, and predicted DNA-DNA melting temperature (T_m_) profiles were calculated using EMBOSS dan v6.6.0 with a window size of 9 nucleotides, a window shift of 1 nucleotide and default DNA and salt concentrations. Stratification of piRNA loci was carried out by calculating the strength of the T_m_ profile as the positive difference of Tm values to background at the high Tm region (TSS to +19), plus the negative difference of T_m_ values to background at the low T_m_ region (+20 to +40). Background was defined as the average Tm at −200 to −100 and +100 to +200 from the TSS.

Quantification of Tm profiles across genomes is challenging given the differences in GC across genomes that result in differences in background nucleotide composition around piRNAs. We calculated this metric in two ways: first, we used a Bayesian Integral Log-Odds approach (BILD) (Yu et al., 2015), using the genome-wide nucleotide composition as background model. This analysis gives a measure of entropy of the sequence at each position, and captures the piRNA termination region as a peak in entropy. We calculated the strength of termination as the area of this peak. Second, we calculated the difference between the maximum Tm value of the high GC content region and the minimum Tm value of the low GC content region.

To compare expression of piRNAs from loci with different pausing signature strengths we used two independent methods to normalize library size. We first normalised piRNA counts to total non-structural mapped reads (non-rRNA and non-tRNA). Second, we fitted a linear model to miRNA counts across pairs of libraries, and inferred size factors as the estimated value of the slope +/-1.96 times the standard error of the estimate.

For analysis of piRNA precursors, reads with starts mapping exactly 2 nt upstream of annotated 21Us were selected. For analysis of TFIIS-derived 3′ cleavage products fragments, all 21U reads were removed to avoid any potential interference from 21U reads coming from annotated and unannotated piRNA loci. Uniquely mapping reads longer than 10 bases mapping at positions +28 to +38 to the TSS were counted in each replicate and normalised to total non-structural mapped reads. The total number of unique sequences was quantified and normalised to the total number of unique non-structural mapped reads. We tested the difference in piRNA precursor length at CIP-RPPH libraries by a Wilcoxon rank sum test.

We used the length distributions of rRNA degradation products (reads mapping sense to rRNA) as a control to rule out variability across libraries due to size selection during library preparation. To compare the lengths of N precursors from library 1 and M precursors from library 2, we sampled N and M rRNA structural reads longer than 23 bases from each library, and calculated a Wilcoxon rank sum test p-value. We repeated the procedure 1000 times to calculate a null distribution of p-values. We noted that differences in sequencing depth can bias the length distribution of unique piRNA precursor sequences, since longer precursors are less abundant. To control for this, we generated 10000 subsamples of 2500 precursor sequences from N2-TFIIS library pairs (3 biological replicates), by weighted sampling of piRNA precursors according to their abundance (number of reads). We generated a distribution of p-values by a one-tailed Wilcoxon rank sum test (TFIIS>N2) and compared this distribution to a null distribution of p-values generated by comparing pairs of subsets of precursors sampled from the N2 library for each biological replicate.

We analysed the parn-1 (tm869) mutant data as described above, with exception of a sample size increase to 5000 piRNA precursor sequences, due to increased sequencing depth in these samples resulting in the detection of a higher number of unique precursor sequences.

## Data availability

Small RNA sequencing data has been submitted to the SRA Study SRP117954: Evolution of piRNAs in nematodes

The *P. oxycercus* and *P. sambesii* genomes have been uploaded to http://caenorhabditis.org/.

## Supplementary Figure Legends

**Figure S1.**
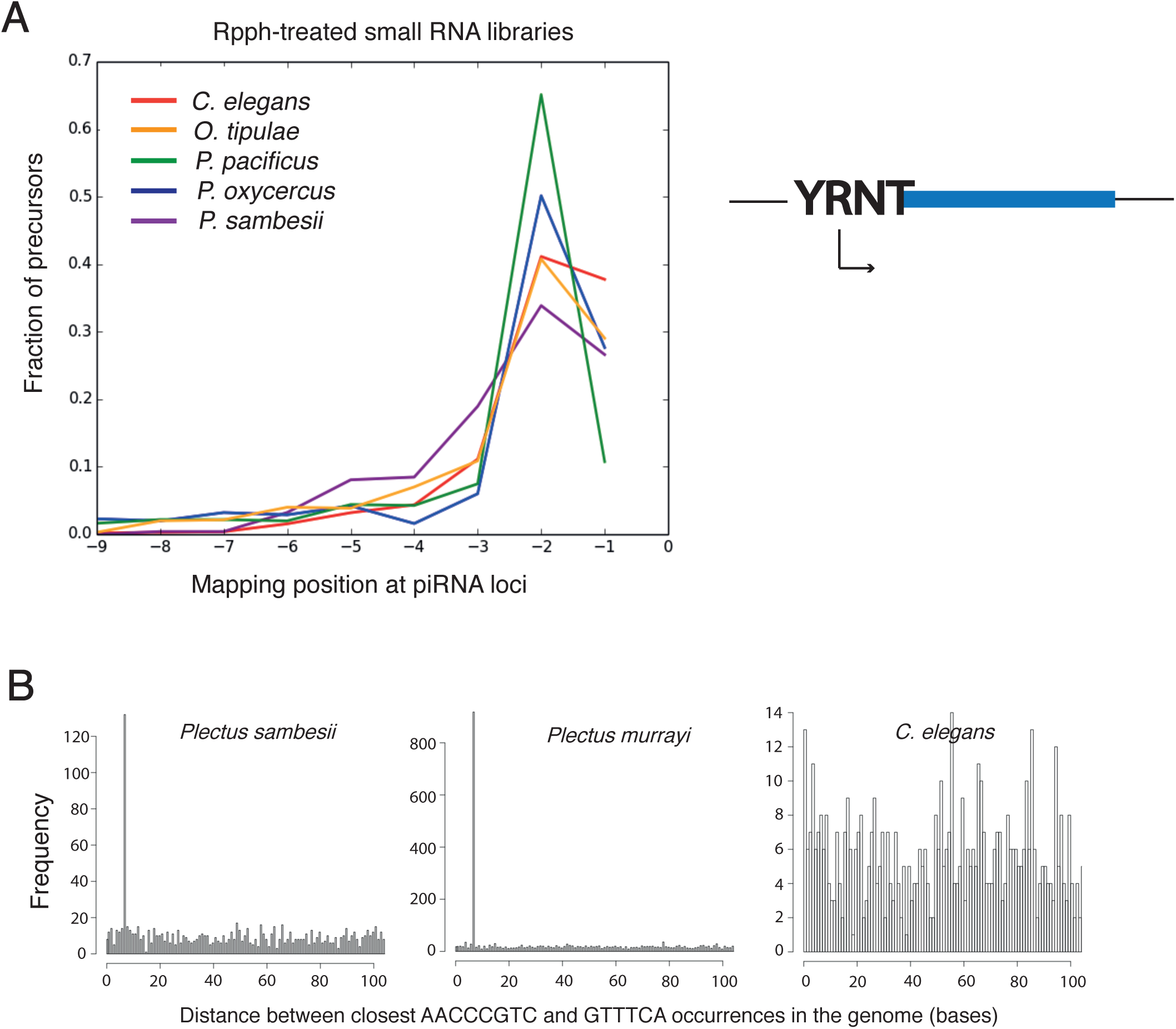
Further analysis of piRNA transcription (relates to Figure 1) A. Transcription initiation at piRNA loci relative to mature 21U sequences across nematodes. 5′ extensions of mapped piRNA precursor transcripts are shown. B. Co-occurrence of GTTTC and AACCCGTC motifs in *Plectus sambesii* and *Plectus murrayi* but not in *C. elegans*. C. *Plectus sambesii* piRNA motifs and SNPC-4 binding sites upstream of U6 snRNAs are the same element. Conserved positions of this motif with the *C. elegans* SNPC-4 binding motif are marked with asterisks.

**Figure S2-1.**
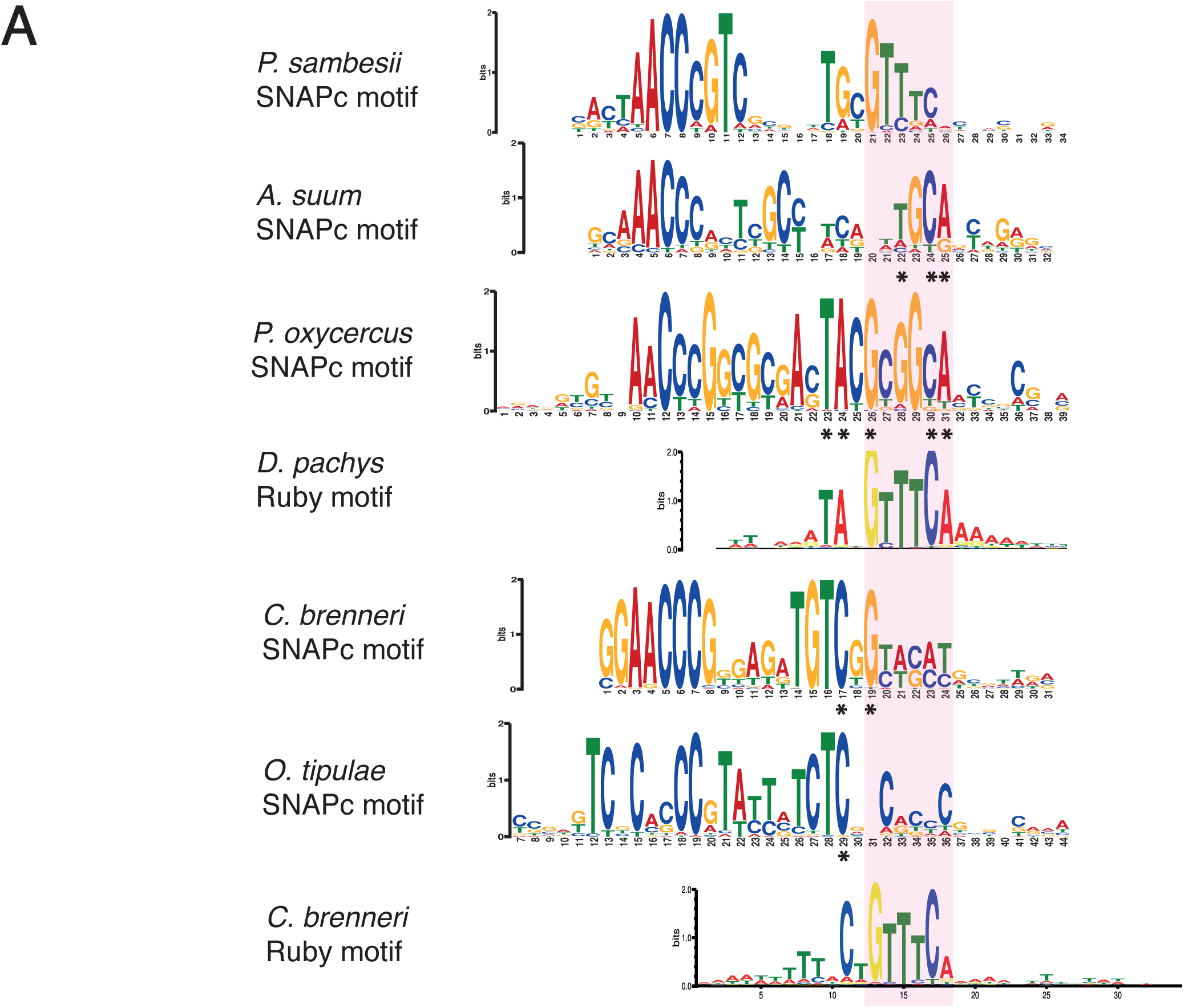
SNAPc and Ruby motif evolution across nematodes (Relates to Figure 2) **A.** Sequence similarities between the 3′ SNAPc and Ruby motifs in several nematode species.

**Figure S2-2.**
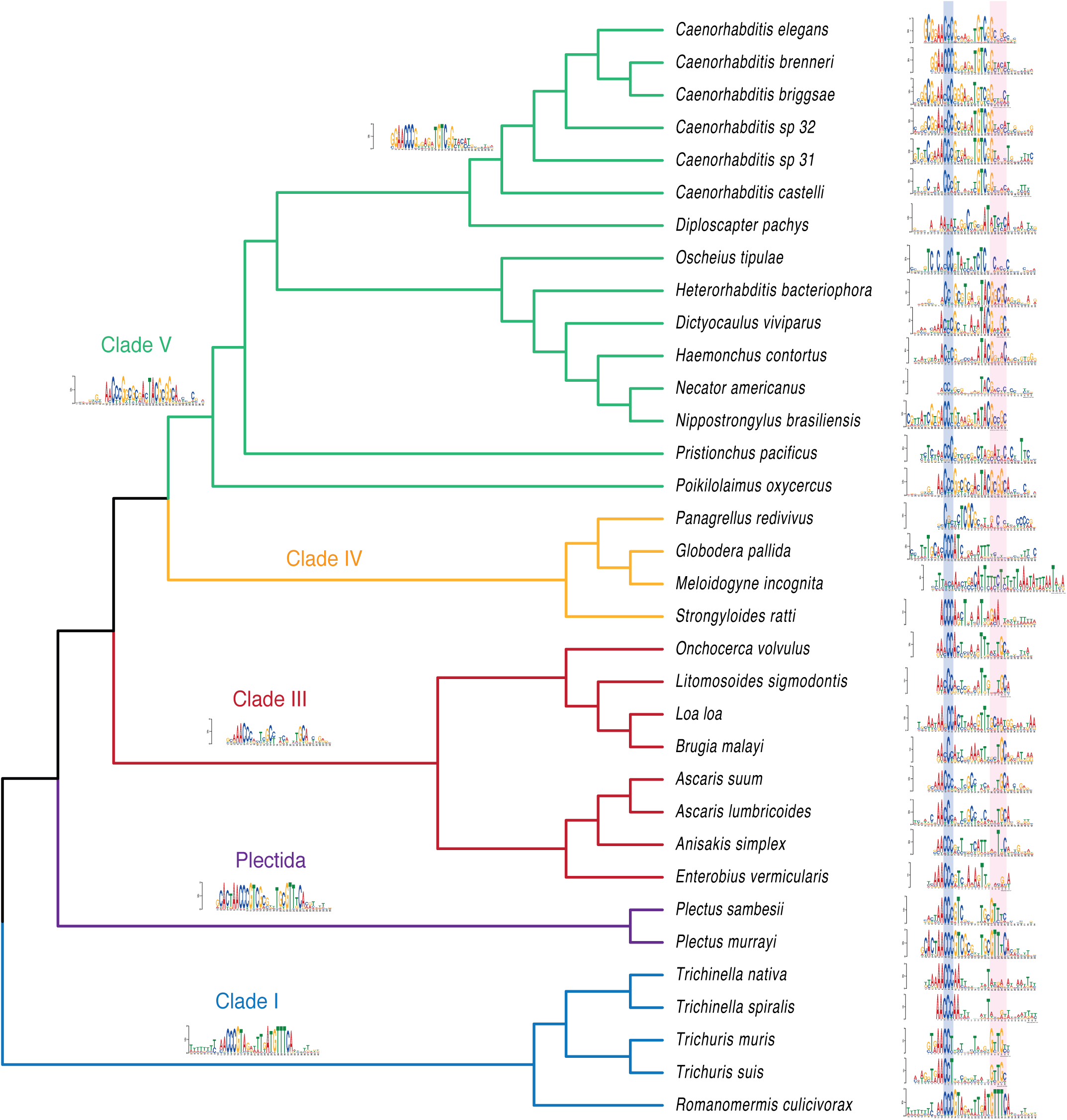
Full tree showing the evolution of the SNAPc motif in nematodes (Relates to Figure 2)

**Figure S3.**
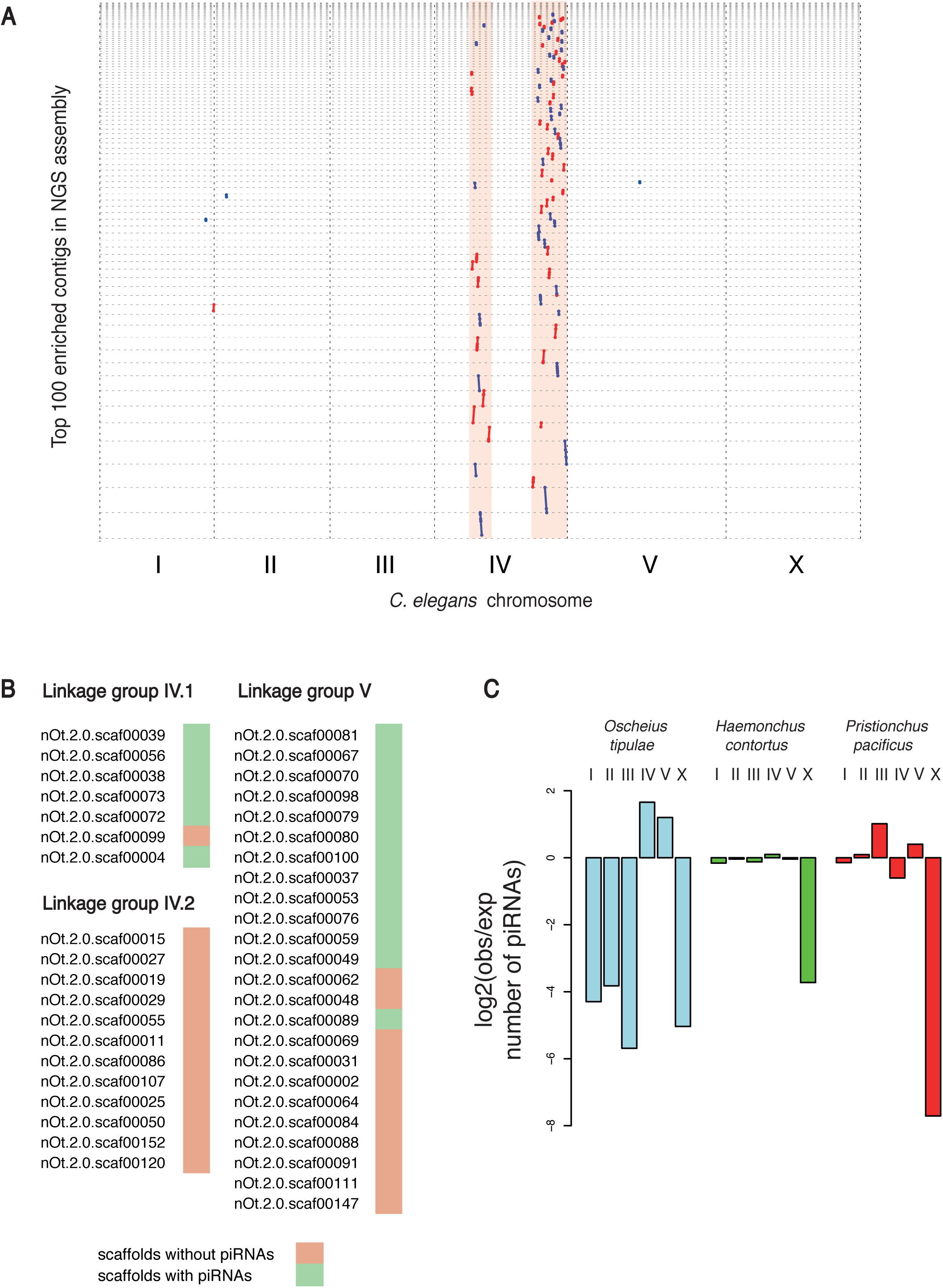
Validation of piRNA clustering analysis methods (Relates to Figure 3) A. Dot plot showing the alignment of the top 100 contigs enriched in piRNAs in a short-read *C. elegans* genome assembly (see Supplementary tables, *Data sources*) to the *C. elegans* reference genome. These map exclusively to *C. elegans* piRNA clusters in IV. B.Identification of scaffolds enriched in piRNAs in the *Oscheius tipulae* linkage map. C.Chromosomal enrichment of piRNAs in *O. tipulae, P. pacificus and H. contortus.*

**Figure S4.**
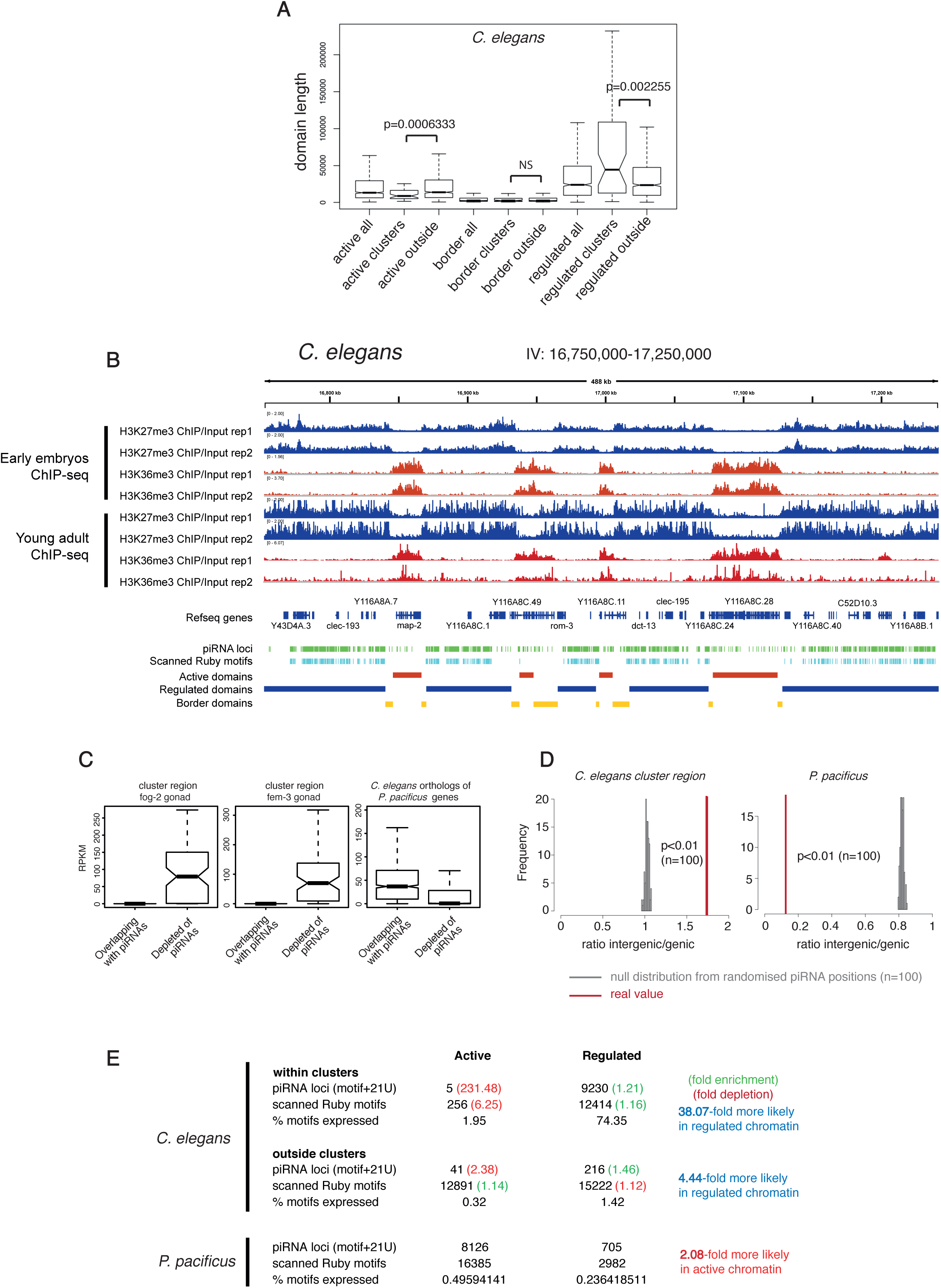
Regulation of piRNA biogenesis by the chromatin environment (Relates to Figure 4) A. Size of regulated and active domains within *C. elegans* piRNA regions compared to the rest of the genome. B. Examples of H3K27me3 and H3K36me3 profiles at piRNA cluster regions in the early embryo and the young adult stage. C. Germline gene expression of genes containing piRNAs in *C. elegans* or the *C. elegans* orthologues of *P. pacificus* genes containing piRNAs. D. Comparison of intergenic/genic piRNA proportions to simulated positions. E. Summary table of annotated piRNA loci and scanned Ruby motifs and their chromatin domain localisations in *C. elegans* and *P. pacificus.*

**Figure S5.**
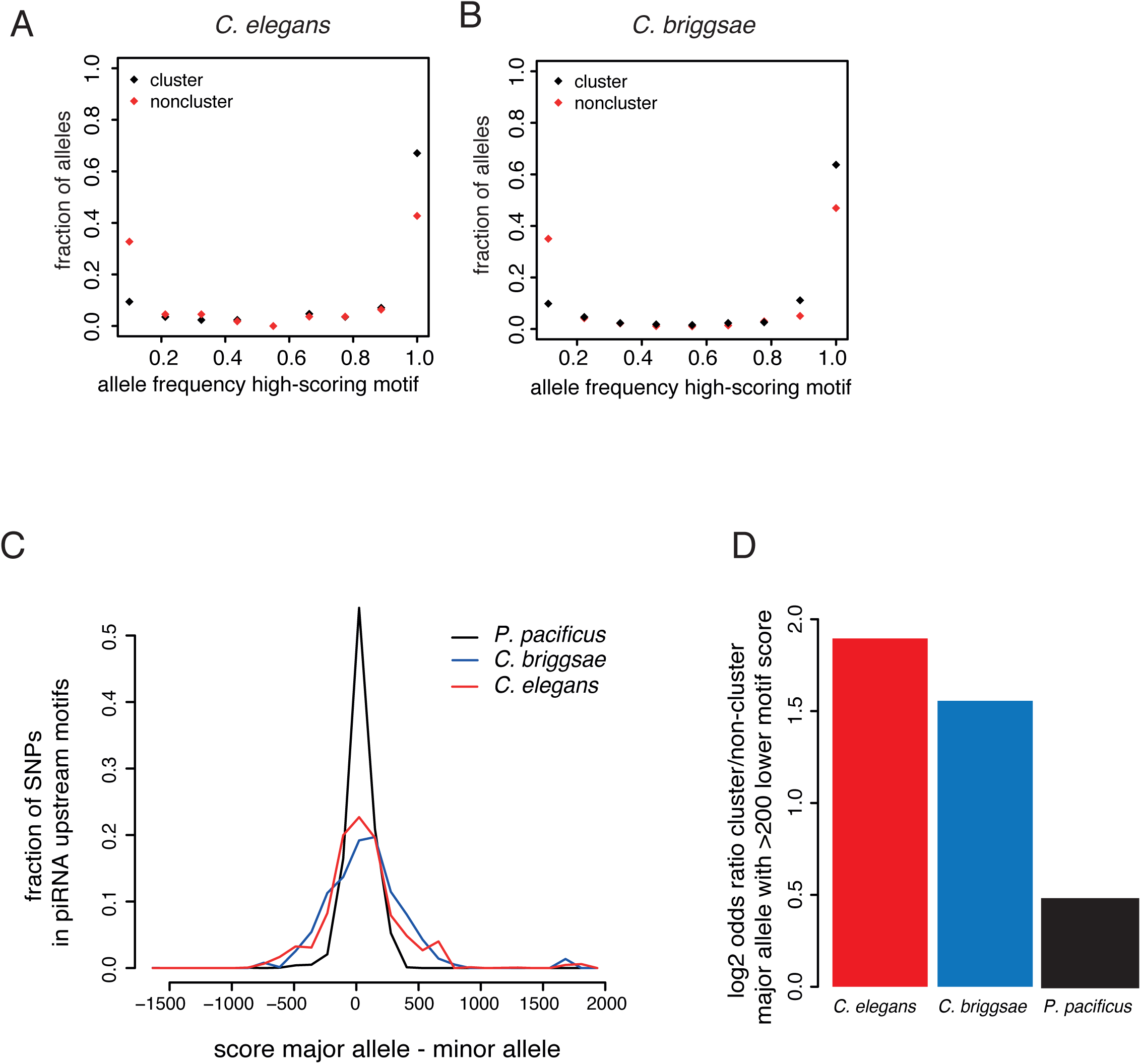
piRNA loci are under selection in both C-type and P-type species (Relates to Figure 5) A-B. Allele frequency distribution of SNPs predicted to alter piRNA motif strength by >400 in *C. briggsae* and *C. elegans*. C. Distribution of the predicted changes in motif scores as a result of all SNPs mapping to piRNA loci in *C. briggsae, C. elegans* and *P. pacificus.* D. Enrichment of high scoring motifs within piRNA clusters in *C. elegans* and *C. briggsae* using the lower threshold of a >200 change in strength, compared to the difference observed between the 10 highest density contigs and the 10 lowest density contigs in *P. pacificus.*

**Figure S6.**
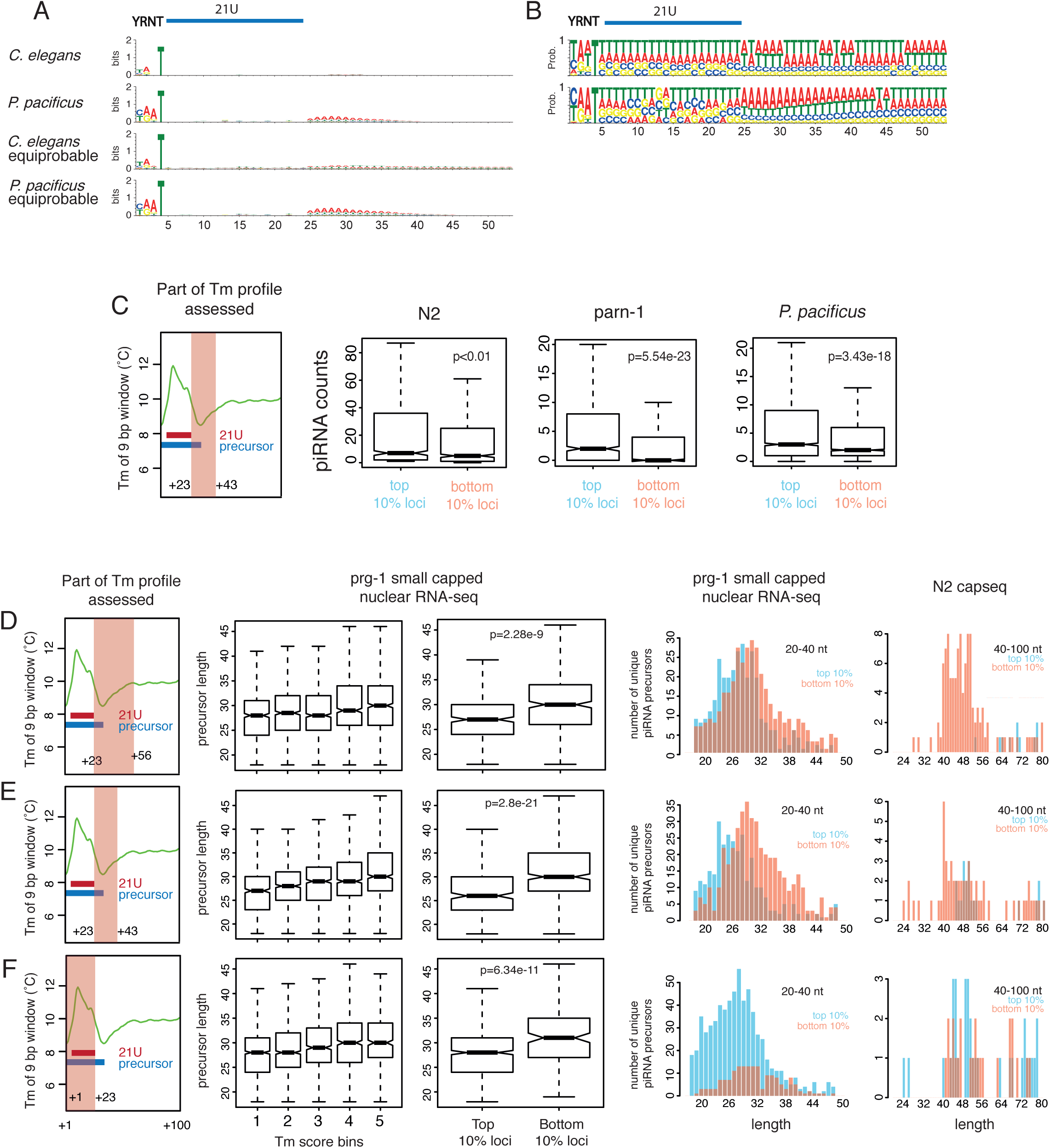
RNA Pol II pausing signals are important for piRNA biogenesis (Relates to Figure 6) A. Entropy sequence logos of piRNA loci in *C. elegans* and *P. pacificus* using either genomic GC content or equiprobable nucleotide frequency as the background model. B. Nucleotide frequency logos of piRNA loci in *C. elegans* and *P. pacificus.* C. Effect of the pausing signal downstream of the 21U sequence on piRNA abundance. D-F. Dissection of the effect of different parts of the pausing-associated sequence signal on piRNA precursor length and abundance.

**Figure S7-1.**
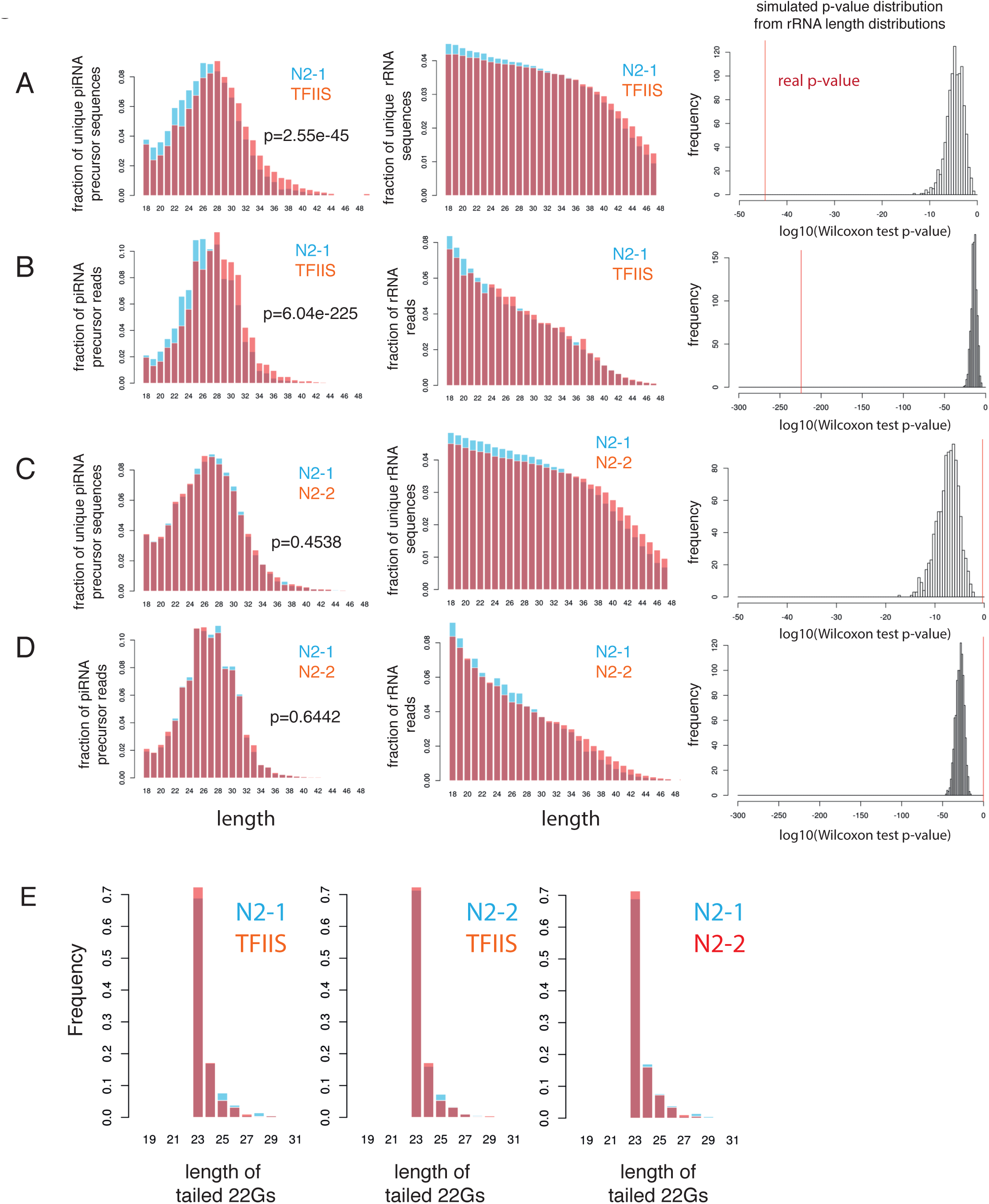
Loss of function of TFIIS modulates piRNA biogenesis in *C. elegans* (Relates to Figure 7) A-D. Comparison of unique and total piRNA precursor reads length distributions in N2 and TFIIS mutant *C. elegans.* The length distribution of rRNA degradation fragment unique and total reads was plotted as a control. P-values for the increase in length were calculated using a Wilcoxon rank sum test, and were compared against a null distribution of p-values calculated by sampling sets of a matching number of unique or total reads from the rRNA length distributions (sampling reads >23 nt). In TFIIS-N2 comparisons, the real p-value is much lower than the calculated null distribution, while in comparisons between two independent N2 libraries, the real p-value is greater, showing that the differences in precursor length are not a result of a global shift in read length in the libraries due to technical variation. E. Length distribution of tailed 22G-RNAs in N2 and TFIIS mutants.

**Figure S7-2.**
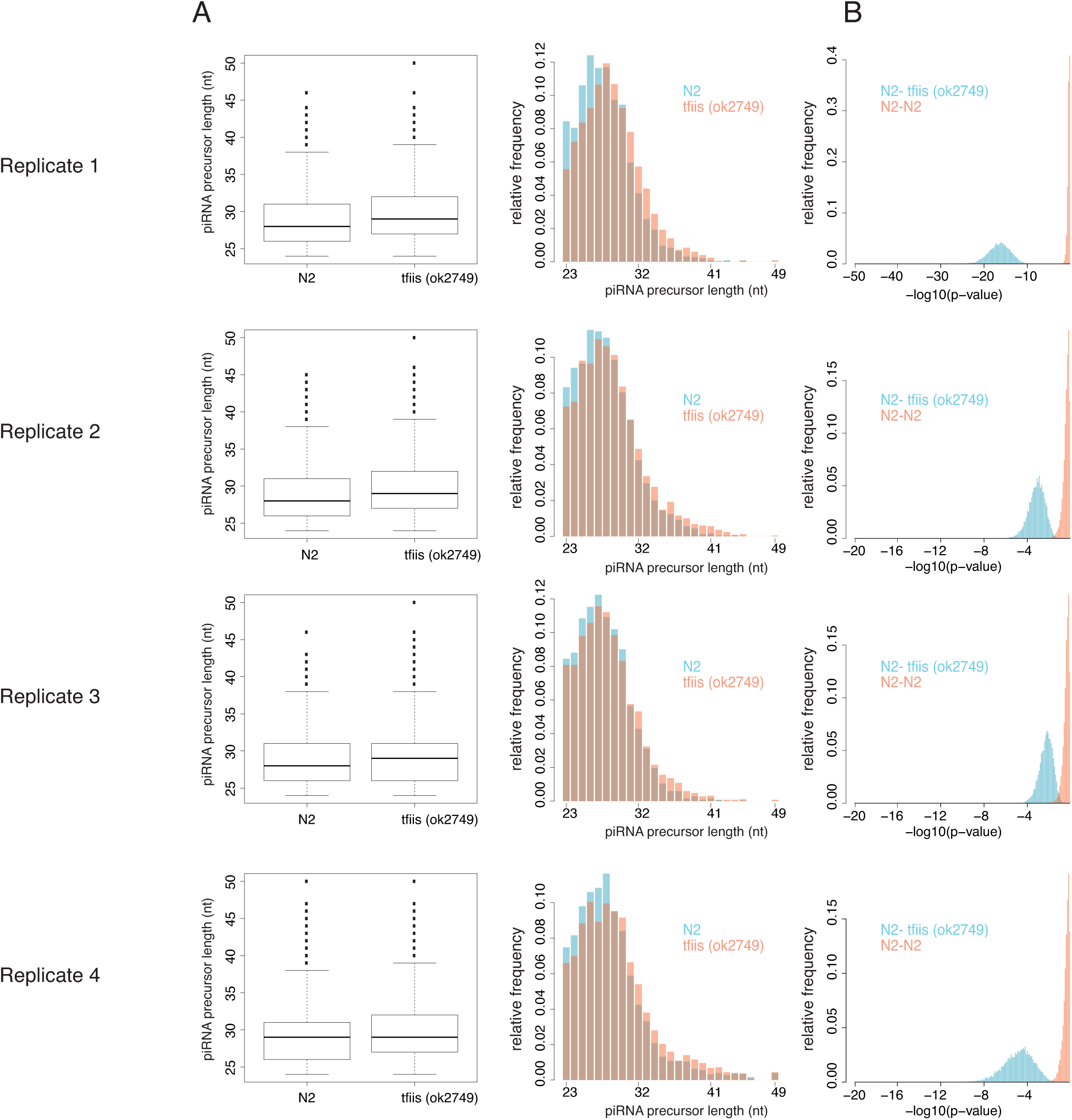
Further analysis of piRNAs in TFIIS mutants controlling for sequencing depth (Relates to Figure 7) A. Length distribution of piRNA precursors in subsets of 2500 precursors sampled according to relative abundance in N2 and tfiis(ok2749) mutants. B. P-value distributions of a Wilcoxon rank sum test for a difference in length, in 10,000 N2 vs TFIIS random subsets (blue) and N2-N2 random subsets (pink).

**Figure S7-3.**
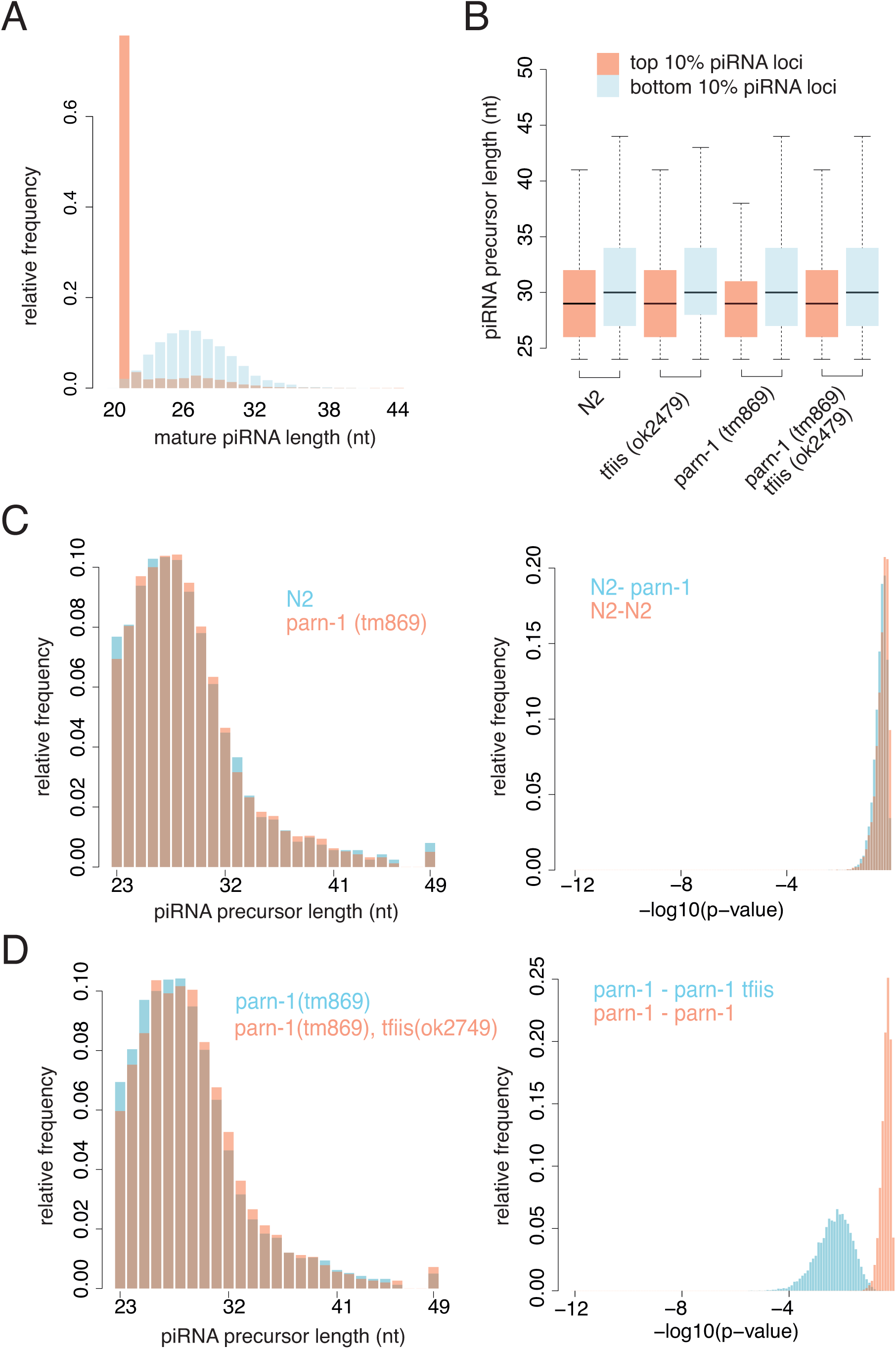
Analysis of piRNA precursors in the absence of trimming. A. Length distribution of piRNA precursors in subsets of 5000 precursors sampled according to relative abundance in N2 compared to parn-1(tm869) mutants. B. piRNA precursor length distributions comparing the top and bottom 10% of piRNA loci stratified according to the strength of the pausing-associated sequence signature. C. Length distribution of piRNA precursors in subsets of 5000 precursors sampled according to relative abundance in parn-1(tm869) mutants compared to parn-1(tm869) tfiis(ok2749) double mutants.

**Figure S7-4.**
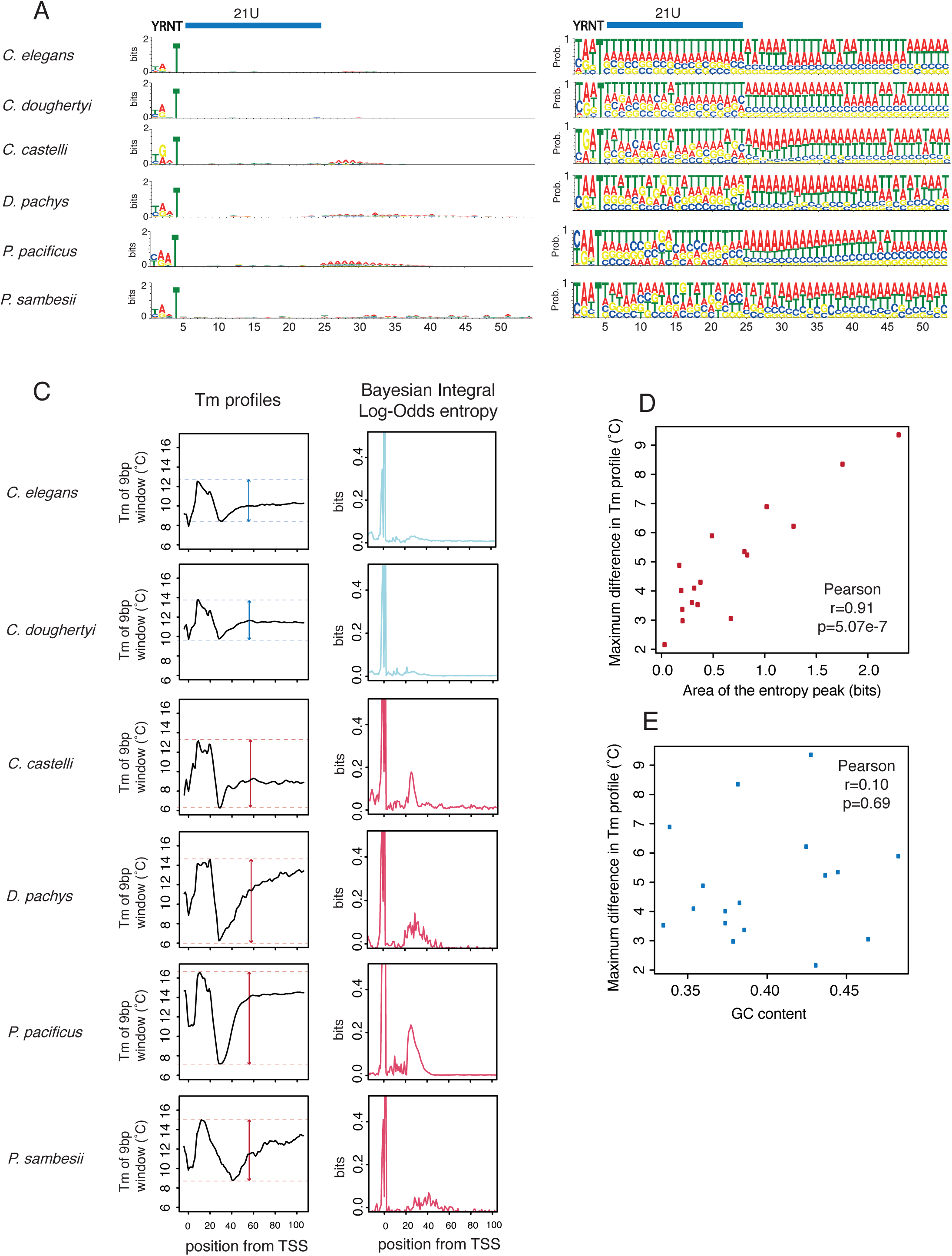
Entropy measurement of the strength of RNA Pol II pausing-associated sequence signatures across nematodes (Relates to Figure 8) A-B. Sequence logos downstream of piRNA loci in several nematodes. C. Melting temperature profiles and Bayesian Integral Log-Odds (BILD) entropy of the sequence around piRNA loci across nematodes. The strength of the pausing signature was calculated as the difference between the maximum value of the high GC content peak and the lowest value of the high AT content region from the Tm profiles, and alternatively as the area under the peak downstream of the 21U relative to background from the entropy profiles. D. Relationship between the different entropy measures used (Spearman r > 0.9). E. Relationship between entropy measures used and GC content (Spearman r <0.1).

## Supplementary tables

Table S1. Data sources

Table S2. piRNA organisation across nematode genomes

Table S3. piRNAs and chromatin domains in *C. elegans* and *P. pacificus*

Table S4. Mature piRNA abundance and pausing-associated sequence signature scores at piRNA loci.

